# Songs show strong individual-level variation and weak population-level dialects in a tropical songbird with genetic structure

**DOI:** 10.1101/2025.11.02.684965

**Authors:** Chiti Arvind, Suyash Sawant, Rohith Srinivasan, G.P. Arpitha, Naman Goyal, Ashwin Warudkar, V. V. Robin

**Affiliations:** Department of Biology, Indian Institute of Science Education and Research Tirupati, Tirupati 517619, India; Department of Wildlife Ecology and Conservation, University of Florida, Gainesville, FL 32603, U.S.A; School of Natural Resources and Environment, University of Florida, Gainesville, FL 32603, U.S.A; Department of Biological Sciences, University of the Pacific, Stockton, CA, 95211, U.S.A; Department of Biological Sciences, Auburn University, Auburn, AL 36849, U.S.A; Section for Macroecology, Evolution, and Climate, Globe Institute, University of Copenhagen, Copenhagen - 2100, Denmark; National Centre for Biological Sciences, TIFR, GKVK Campus, Bangalore 560065, India

**Keywords:** Birdsong syntax, population genetics, vocal learning, sky island, landscape ecology

## Abstract

Dialects in birdsongs represent population clusters and cultural diversity. Cultural and genetic variation in animals, however, do not always evolve along the same trajectory. Few studies have tested this in a wild songbird with a continuous learning strategy. Songbirds (oscine Passeriformes) with a continual learning window exhibit great song diversity and may show cultural differentiation that outpaces genetic divergence. We investigate cultural and genetic variation in a tropical endemic songbird in a topographically and climatically complex sky-island landscape of the Western Ghats of India. We combine fine-scale acoustic, behavioural, and genetic data: from 90 individuals (10,816 songs), analysed using frequency-temporal parameters and a linguistic method for syntax; a playback experiment with 136 trials to compare behavioural responses to local and non-local songs; and blood samples from 242 individuals (10,638 SNPs) to infer genetic population structure. We find no difference in songs across locations and individuals in the time-frequency domain, but only in syntax. One-third of notes, the smallest vocal units, are widely shared across all individuals in their distribution. Longer note combinations result in greater individual differences within populations, but only form small differences between populations. Geographic distance strongly predicts genetic but not song syntactic differences, and playback experiments show equal responses to local and non-local songs. Thus, strong individual signatures but weak local-level dialects are generated by a shared note repertoire and associated combinatorial syntax. We highlight that cultural transmission may operate at fine social scales, decoupled from genetic structure, which is strongly driven by isolation-by-distance.

**Significance statement:** Whether learned behaviours exhibit patterns similar to those of inherited genes remains a long-standing question in evolutionary biology. Birdsong has a strong cultural component paralleling human language, yet most knowledge comes from birds with a narrow learning window. Open-ended learners, who acquire new songs throughout life like humans, are poorly documented in the wild. We studied such a songbird restricted to the tropical cloud montane forests of southern India. Songs appear acoustically similar across populations but differ in how basic units are ordered (syntax), while individuals are more distinct from each other in longer note combinations than populations are. Genetic similarity decreased with geographic distance. Culturally learned traits can thus evolve independently of genetic structure, showing how learning decouples behaviour from geography-driven genetic ancestry.

## Introduction

A species may exhibit different patterns of spatial variation in cultural and genetic traits due to its mode of inheritance. Genetic information is passed down through generations, while culturally transmitted behaviours are learned from related or unrelated conspecifics(Cavalli-Sforza & Feldman, 1981; Whiten, 2021) and can form local traditions(Schuppli & van Schaik, 2019; Whiten, 2019). The relationship between cultural and genetic transmission is of interest, as behavioural traits (such as vocalisations) under sexual selection can also contribute to genetic divergence(Beans, 2017; Slabbekoorn & Smith, 2002; Whitehead et al., 2019). Birdsong, often a culturally transmitted trait(L. Aplin, 2022; L. M. Aplin, 2019), can often have dialects(Podos & Warren, 2007). However, patterns of differentiation across cultural and genetic axes in populations might not follow the same trend(Luo et al., 2024; Nwankwo et al., 2018).

Initial work on cultural exchange and gene flow suggests that genetic distances result from cultural dialect boundaries acting as premating isolation barriers via the genetic adaptation hypothesis(Baker, 1975; Baker et al., 1982; Baker & Mewaldt, 1978). However, after accounting for spatial autocorrelation, it was found that genetic distances were explained by geography rather than dialect boundaries ((Zink & Barrowclough, 1984). In line with this, genetic distances following an isolation-by-distance pattern are supported by other studies and highlight that populations with distinct dialects do not always form genetic barriers(MacDougall-Shackleton & MacDougall-Shackleton, 2001), 2021; Soha et al., 2004; Poesel et al., 2017). Such discordance between cultural and genetic variation is driven by different rates of evolution: cultural turnover occurs more rapidly over a short period (a single generation) than genetic signals.

Birdsong playback experiments serve as a key tool in testing cultural differences across genetically dissimilar populations. Birds usually respond more strongly to local songs rather than those from non-local populations(Colombelli-Négrel & Kleindorfer, 2021; Lipshutz et al., 2017; Luo et al., 2024; Sebastián-González & Hart, 2017). However, several songbirds also exhibit individual variation ((Sandoval et al., 2014);(Průchová et al., 2017)(Petrusková et al., 2016)), which is often less than the variation across widely separated populations(Węgrzyn et al., 2025). In some cases, although rarely documented, this individual variation can mask geographic song variation even in geographically separated populations(Oñate-Casado et al., 2026).

Cultural variation in songbird vocalisations also depends on learning strategy. Closed-ended learners develop fixed, crystallised songs, whereas open-ended learners retain continual learning windows and greater vocal variability(Beecher & Brenowitz, 2005). Much of the theory of how bird song variation comes from closed-ended learners, largely in Europe and the Americas. Studies on song variation in those with lifelong learning windows remain limited(Sprau & Mundry, 2010; Vokurková et al., 2013; Wheatcroft et al., 2022) despite the parallels they draw towards continual vocal learning in human languages in terms of song learning neural pathways(Hyland Bruno et al., 2021) and hierarchical structure(Berwick et al., 2011). Whether geographic distance predicts song differentiation in open-ended learners remains unresolved. Cultural variation due to geographical distances is seen across continental scales in African red-faced warbler *Cisticola erythrops*(Benedict & Bowie, 2009) or at finer scales in Great Reed warblers *Acrocephalus arundinaceus* (Węgrzyn et al., 2025), the latter of which also exhibits population-level dialects at shorter distances (<50km) (Węgrzyn et al., 2025). However, testing for behavioural discrimination in open-ended learners remains limited. Playback experiments with an open-ended learner that produce complex vocalisations, such as those of the pied flycatcher (*Ficedula hypoleuca*), indicate that individuals respond more strongly to local than to foreign calls(Gallego-Abenza et al., 2025).

Apart from geographic patterns, the song’s structure often provides additional resolution. In species with complex songs, the vocal syntax, i.e. the sequential arrangement of syllables drawn from a shared set of basic unit types, can differentiate individuals and populations more precisely than spectral features (measurements of frequency and time) alone (Marler and Pickert, 1984; Lachlan et al., 2014; Sawant et al, 2025; Zsebok et al., 2021;(Berwick et al., 2011; Kipper et al., 2004; Kroodsma, 1980; Mota & Cardoso, 2001; Petrusková et al., 2010; Sawant et al., 2025).

Beyond song-learning pathways, vocal cultural variation is also shaped by isolation-by-distance, resistance, and environment, and these barriers may also differentially affect genetic variation. Isolation-by-distance patterns of song dialects and genetic population structure could be concordant, where geographically distant populations are genetically and vocally distinct(Camacho-Alpízar et al., 2018; Danner et al., 2011, 2017; Luo et al., 2024) or discordant, where there is high gene flow between populations, yet populations exhibit distinct vocal dialects(Poesel et al., 2017; Searfoss et al., 2020). Physical barriers or habitat heterogeneity (natural or anthropogenic) can affect movement at various scales(Ceresa et al., 2023; Hindley et al., 2018; Musher et al., 2022), resulting in an isolation-by-resistance(McRae, 2006). Lastly, climatic variation can result in an isolation-by-environment, which can drive song and genetic differentiation at fine scales(Moore et al., 2005; Quintero et al., 2014). Considering these modes of isolation, we need more integrative approaches combining behavioural ecology and landscape genetics to assess the drivers of cultural and genetic variation within a species.

We examine song and genetic variation in the white-bellied sholakili (*Sholicola albiventris*), an open-ended learner(Sawant et al., 2022, 2025) with annual song turnover across individuals (Sawant et al., 2025). *S. albiventris* is a non-migratory, year-round territorial bird, endemic to the Shola Sky Islands in the Western Ghats of India, with over two decades of research(Arunima et al., 2026; Robin & Sukumar, 2002). Within a Sky Island, there is an east-west cline and asynchrony in precipitation (wetter in the west, with higher rainfall earlier in the year), which has been linked to genetic differentiation in many taxa, such as frogs(Janani et al., n.d.) and butterflies(Sekar & Karanth, 2013). The landscape is patchy (see below for further details). Thus, *S. albiventris* in its natural habitat provides a system to assess cultural and genetic variation across modes of isolation via distance, environment, and landscape resistance.

Previous work on *S. albiventris* has revealed both spectro-temporal and syntactic differences across sky islands, as well as between locations at the extreme ends (60 km apart) within a single island(Purushotham & Robin, 2016; Robin et al., 2015). These cultural differences have been paired with limited contemporary gene flow using microsatellite loci, suggesting effects of anthropogenic land-use modification on habitat change(Purushotham & Robin, 2016; Robin et al., 2015). Over the last half-century, an increase in invasive timber plantations has modified the naturally biphasic landscape (of shola grassland and shola forest) and increased overall woodland cover(Arasumani et al., 2019). This modification has affected species’ occupancy(Jobin et al., 2025; Lele et al., 2020) and provides a unique configuration of increased connectivity(Arunima et al., 2026) in the east and a stable historical connectivity in the west. Given the species’ ecology, including its song, landscape characteristics may interact to affect genetic and cultural connectivity. One could expect greater cultural connectivity from the increased connectivity, relative to genetic differences that take time to accumulate(Whitehead, 2017).

In this system, we a) assess patterns of song sharing (using frequency-time parameters and syntax) across the species distribution, b) test behavioural discrimination towards songs from local vs non-local populations using playback experiments, c) examine patterns of genetic population structure using Next Generation Sequencing techniques, and d) infer concordance or divergence in song variation and genetic population structure considering isolation barriers in a landscape ecology framework. With detailed knowledge of the species(Purushotham & Robin, 2016; Robin et al., 2011, 2015, 2017; Sawant et al., 2025), we were able to design our sampling and expect that overall song sharing will decrease with distance, with individuals within the same location sharing more songs. Regarding behavioural responses, birds are expected to respond more strongly to local songs than to non-local songs, reflecting a cultural affinity for local songs. We expect some admixture in the genetic population structure, highlighting an isolation-by-distance pattern, and perhaps an impact of the climatic barrier. In areas with recent land-use changes and increased connectivity (Arunima et al., 2026), we also expect an increase in song sharing (in the east) compared to native patchiness (in the west). A discordance in the song variation and genetic structure is expected, as cultural changes in vocal behaviour may accumulate faster than genetic differentiation.

## Methods

### 2.0 ) Study site and field sampling

We studied *S. albiventris* individuals (Figure 1a) from 13 locations across two Shola Sky Islands (Palani-Anamalais and Highwavies), which are 1400 m above sea level and connected by a 1,000 m ridge (Figure 1b). This landscape comprises a natural mosaic of shola forest and shola grasslands (Robin & Nandini, 2012), interspersed with human-modified stretches, including tea and timber stands (such as those of *Acacia, Eucalyptus,* and *Pinus* - Arasumani et al., 2018) (Figure 1c). This habitat heterogeneity leads to varying levels of isolation, reducing connectivity among *S. albiventris* individuals (Figure 1d). An asynchronous pattern of precipitation across the landscape over a short 60km also supports environmental variation (Figure 1e). The sampled locations were approximately 10km apart, except for the stretch between Mathikettan (in the Anamalai-Palanis) and Highwavies (the southernmost location), where the elevation was lower (a 1000m ridge), and the population was too small to sample.

**Figure 1:**
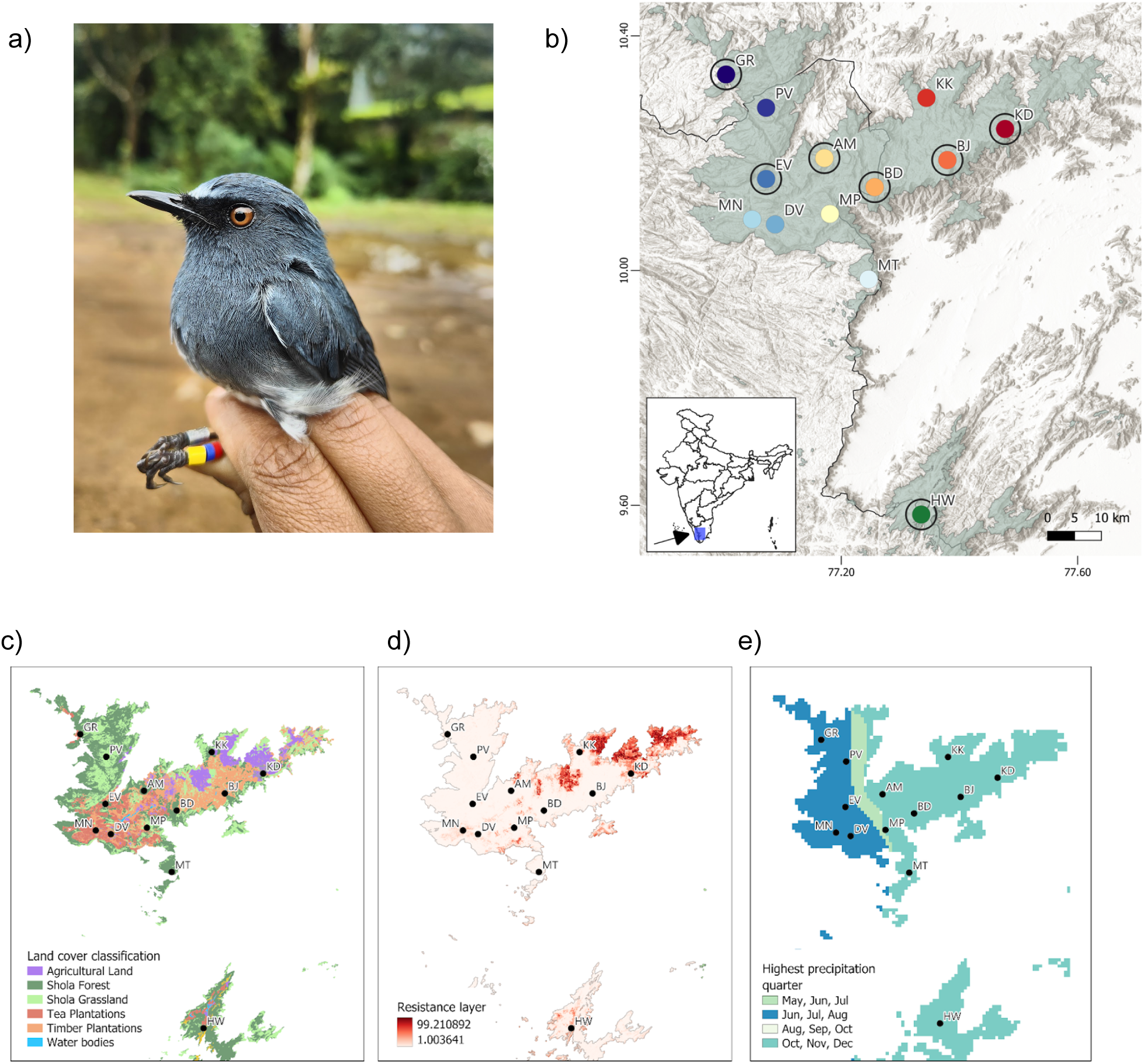
a) A colour-banded individual of S. albiventris **b)** Sampling locations: All coloured dots represent locations sampled, with genetic data (N = 13) (Grasshills - GR, Poovar - PV, Munnar (MN), Eravikulam (EV), Anaimudi (AM), Meeshapulimala (MP), Mathiketten (MT), Bandar (BD), Kookal (KK), Berijam (BJ), Kodaikanal (KD), Highwavies (HW), with warm to cool colours along an east-to-west gradient, and green is the southernmost location. The solid outer circles are locations sampled for songs (N = 7). Playback experiments were conducted at one location: KD. The pale green-shaded area indicates the 1400m elevation contour encompassing the Annamalai-Palani plateau (north) and the Highways range (south); the base layer represents terrain contours. The solid black line represents the state boundary of Kerala (left) and Tamil Nadu (right). The inset map shows peninsular India with a blue rectangle highlighting the Anamalai-Palani and Highwavies ranges. **b)** land cover classification showing both natural and anthropogenic landscape heterogeneity (Arasumani et al, 2018), **c)** resistance layer with lighter colour showing low resistance and dark colours showing higher resistance (Arunima et al, 2025), **d)** asynchrony in precipitation with the highest rainfall quarter highlighted in the landscape.

Genetic data collection: We captured 332 birds (292 from 2022-2024; 40 from 2012-2018) using mist nets, banded them with a site-unique colour band and a unique alpha-numeric metal band (Supp. Table 1, Figure 1a). We collected a few drops of blood samples from all individuals following Robin et al. (2010), and blood was stored in Queens Lysis Buffer and on FTA cards. All individuals were molecularly sexed as male or female using P2/P8 markers (Griffiths et al. 1998), since these birds are monomorphic. For genomic analyses, we used samples from both sexes and restricted samples per location to at least 10 individuals, totalling 242 (22 samples from 2011 - Robin et al., 2015) (Supp. Table 1).

**Table 1:** Linear Mixed model output for vocal behavioural responses (PC1) to local and non-local stimuli, considering the local population as the reference. Estimates along with 95% Ci are shown.

| PC1 ~ pb_loc |  |  |  |
| --- | --- | --- | --- |
| Predictors | Estimates | CI | p |
| (Intercept) | 0.20 | -0.33 – 0.72 | 0.461 |
| pb loc [non-local_near] | -0.38 | -0.81 – 0.05 | 0.080 |
| pb loc [non-local_far_South] | -0.35 | -0.79 – 0.09 | 0.117 |
| pb loc [control] | -1.67 | -2.26 – -1.08 | <0.001 |
| **Random Effects** |  |  |  |
| $\sigma^2$ | 0.85 | | |
| $\tau_{00}$ focal_indv | 0.72 | | |
| $\tau_{00}$ observer | 0.07 | | |
| ICC | 0.48 |  |  |
| N focal_indv | 28 |  |  |
| N observer | 8 |  |  |
| Observations | 136 |  |  |
| Marginal R<sup>2</sup> / Conditional R<sup>2</sup> | 0.122 / 0.546 |  |  |

Acoustic data collection: Focal handheld recordings of male individuals that were taken from colour-banded and some behaviourally male individuals (unbanded and spatially identified based on site affinity to GPS recording points for a single season) were collected from *S. albiventris* across seven locations to capture the east-to-west gradient and the southernmost population above the Shencottah Gap (Figure 1b). At each location, focal recordings were obtained from 12–14 individuals over 10–14 sampling days during a single breeding season (March–June, 2022–2024). This sampling approach captured immediate song variety at the individual level. Recordings were made using either a parabolic microphone (Wildtronics Pro Mono) or a shotgun microphone (Sennheiser ME 66), paired with a Zoom H4N digital recorder (44.1 kHz sampling rate).

### 2.1 ) Acoustic data analysis

Song analysis was done to assess spectral variables (frequency-time parameters) and syntax. To equally represent all populations (for syntax), we selected a standardised number of 1000 songs per population to ensure equal sampling across populations, totalling 90 individuals (i.e., 12-14 individuals per population) with a minimum of 20 songs per individual, capturing immediate variety (Supp. Table 2).

**Table 2:** Linear Mixed model output for physical behavioural responses (PC2) to local and non-local stimuli, considering the local population as the reference. Estimates along with 95% Ci are shown.

| PC2 ~ pb_loc |  |  |  |
| --- | --- | --- | --- |
| Predictors | Estimates | CI | p |
| (Intercept) | 0.20 | -0.14 – 0.54 | 0.254 |
| pb loc [non-local_near] | -0.18 | -0.62 – 0.27 | 0.430 |
| pb loc [non-local_far_South] | -0.28 | -0.74 – 0.17 | 0.220 |
| pb loc [control] | -0.36 | -0.97 – 0.25 | 0.245 |
| **Random Effects** |  |  |  |
| $\sigma^2$ | 1.01 | | |
| $\tau_{00}$ focal_indv | 0.07 | | |
| ICC | 0.06 |  |  |
| N focal_indv | 28 |  |  |
| Observations | 136 |  |  |
| Marginal R<sup>2</sup> / Conditional R<sup>2</sup> | 0.015 / 0.078 |  |  |

Each note in a song was manually annotated using Raven Pro v1.6 (K. Lisa Yang Centre for Conservation Bioacoustics at the Cornell Lab of Ornithology). A note is defined as the smallest unit of a song (Catchpole & Slater, 2003), and a continuous trace on the spectrogram (Supp. Figure 1, following Sawant et al., 2025) and notes separated by less than 2ms were marked as an instance of a Complex Vocal Mechanism(CVM) (following (Purushotham & Robin, 2016; P. Singh & Price, 2015). This approach yielded a total of 6,997 songs (of good signal-to-noise ratio as visually inspected) and a final pool of 87,589 notes. We then extracted seven spectral parameter variables (listed in Supp. Method S1.1) at the song level and calculated the proportion of CVMs per song (i.e., the instances of CVMs in a song divided by the total number of notes in a song), where a difference across populations is linked to cultural transmission (Purushotham & Robin, 2016; P. Singh & Price, 2015). We then visualised population-level frequency-time parameters and the proportion of CVMs using a Principal Component Analysis (PCA) with the prcomp package ((R Core Team, 2024). We also used a nested random-effects model to partition the variance in song-level frequency-time measurements (log-transformed) into within-individual, within-population between-individual, and between-population components (Lme4 package).In addition, we examined population-level variation in the proportion of Complex Vocal Mechanisms (CVMs) along a west-to-east gradient (Generalised Linear Mixed Models using the glmmTMB package; Brooks et al.,(Brooks et al., 2017)).

Next, to assess syntax, we first denoised notes using a two-step, semi-automated process (custom-modified scripts; Sawant et al. 2025) and then classified them using spectrogram cross-correlation (Supp. Method S1.2). Notes were classified with a 75% accuracy (Supp. Methods S1.3). Subsequently, we derived syntax combinations using a linguistic method of N-grams to group notes (1-grams) into longer note strings in pairs (2-grams) or triples (3-grams) (Sawant et al., 2025). We then retained N-gram types after filtering for their occurrences within each population, resulting in a total of 315 1-grams (occurring more than 5 times), 1789 2-grams (occurring more than 2 times) and 13373-grams (occurring more than 2 times) and assessed song sharing between populations across different levels of syntax (Supp. Method S1.3). We also used a simulation test to assess whether combinatorial syntax (2-grams and 3-grams) in an individual’s songs is non-random. For this, we preserved individual-level note (1-gram) frequencies along with the notes per song sampled; then randomly shuffled the individuals’ note pool to generate 2-gram and 3-gram sequences and compared them to the empirical values.

To quantify differences in N-gram sharing within and among locations across syntax levels for an individual, we conducted a PERMANOVA (999 permutations; vegan) for each N-gram type (1-gram to 3-gram). We used both presence-absence (via Jaccard’s distance) and relative abundances (via Bray–Curtis distance, weighted by the total number of notes per individual), along with calculating effect sizes (omega square, package MicEco, (Russel & Oksanen, 2025). In addition, we calculated within-population-level individual heterogeneity (beta dispersion; function PERMDISP in the vegan package). To test for an increase in individual identity with increasing syntactic complexity (1-gram to 3-gram), we used a generalised linear mixed model regression with the predictor variable as individual dispersion distances from their population centroid (from PERMDISP, for each N-gram level) and response variable as the N-gram length, along with a random effect as an individual nested within a population (package glmTMB).

In addition, to obtain population-level song distances, we computed distances between centroids (NMDS via the vegan package - Oksanen et al., 2024) based on the presence-absence and relative abundance matrices for each N-gram type (1-gram, 2-gram, and 3-gram). These two location-wise song distance matrices (presence-absence and relative abundance) were used to assess the effects of modes of isolation (distance, environment, and resistance) on song differences (section 2.4).

### 2.2 ) Experimentally testing discrimination towards songs from different locations using playback experiments

Playback experiments were conducted at Kodaikanal (KD) using local and non-local male songs. We conducted 136 trials on 28 male birds (11 colour-banded males and 17 unbanded - verified based on pairs with banded females and territory mapping) over two seasons (winter 2024 and summer 2025). A playback trial consisted of a six-minute trial comprising the playback stimulus (3 minutes) followed by three minutes with the speaker turned off. For a trial focal male was subjected to a single playback stimulus (3min) consisting of male songs from one of the three locations which included 1) local songs (KD), 2) non-local near songs (Berijam: BJ, ∼10 km from KD), 3) non-local far songs (Highwavies: HW, >50 km from KD), or a 4) control (soundscape stimulus without *S. albiventris* vocalizations).

All playback stimuli were renamed and kept blind to the observer team, and each playback trial was randomly chosen for a focal individual. Each focal male was presented with two distinct stimuli on a given day (one in the morning, 6 to 8 am, and one in the evening, 4-6 pm), and it was ensured that neighbouring males did not receive the same stimuli. Each location was also represented by three stimuli, comprising song recordings from three different male individuals, to avoid pseudoreplication (following (McGregor, 2000). Vocal responses were recorded using a directional microphone (Sennheiser ME 66) with a Zoom H4n Pro recorder, and annotations were made of vocal and behavioural measures via a double observer system to ensure the focal bird was always in view (Moseley et al., 2013, 2019) (Supp. Methods S2.0).

Four major parameters noted (following (Lipshutz et al., 2017) were: 1) average distance to the speaker for the duration of the experiment, 2) total singing duration of the focal individuals, 3) number of flights over the speaker, and 4) latency to sing. Stronger responses would entail a shorter average distance to the speaker, longer singing duration, more flights over the speaker, and lower latency to sing. The four observed variables were scaled and centred to generate a composite score for behavioural response by performing a Principal Component Analysis (package princomp). To test for differences in behavioural responses to local and non-local (near and far) songs, we used a linear mixed model (package lme4; Bates et al., 2015). The predictor variables included the population from which the playback songs originated, and the random effects included were the observer annotating the experiment and the focal bird.

### 2.3 ) Genomics

We extracted DNA from both sexes using the Qiagen DNeasy Blood and Tissue Kit (Qiagen, Hilden, Germany), following the manufacturer’s protocol with two modifications: an extended incubation period (two hours) and an increased quantity of AL buffer (300 μL). In two sets (159 and 83 samples), we prepared ddRADseq libraries (following Tyagi et al, 2024, Praveen et al., 2024; Supp. Methods 3.0) and sequenced on an Illumina Novaseq 6000 platform. Reads from all runs were merged and processed together. An interactive assembly pipeline, ipyrad v.0.9.105 (Eaton & Overcast, 2020), was used for variant calling with the reference assembly method, by mapping reads to a high-quality reference genome of the *S. albiventris* (Vinay et al., 2025), and we filtered SNPs using vcftools (Supp. Methods 3.1).

To assess historical differentiation across the 13 sampled locations, we calculated Hudson’s FST (Hudson et al., 1992) ( smartpca in EIGENSOFT, Price et al., 2006) that accounts for unequal sample sizes across populations (Bhatia et al., 2013). To quantify contemporary differentiation, we calculated DPS (1 – the proportion of shared alleles; PopGenReport - Gruber & Adamack, 2015). To assess population structure, we used two different approaches: 1) a Principal Component Analysis (package PLINK - Purcell et al., 2007) using Euclidean distances (visualised using ggplot2 in R), and 2) a population classification method, ADMIXTURE (Alexander et al., 2009). We ran ADMIXTURE for 10-fold cross-validation (with K, i.e., population clusters ranging from 1 to 10), selecting the preferred K based on the lowest cross-validation error. Results were visualised using the pophelper package (Francis, 2017) in R. We also assessed genetic co-ancestry among individuals within and across locations (fineRADstructure v.0.1 (Malinsky et al., 2018). To detect barriers to dispersal that may affect isolation-by-distance, we employed FEEMS (Fast Estimation of Effective Migration Surfaces, (Marcus et al., 2021). This technique generates migration surfaces and quantifies pairwise edge estimates between nodes, representing effective migration rates that deviate from expectations under isolation-by-geographical-distance (IBD). Cross-validation was performed across the smoothing parameters lambda (0.001-100, intervals = 20) and lambda_q (0.001-100, intervals = 5), with the optimal lambda selected for the spatial graph based on minimising cross-validation error (Supp. Methods 3.2).

### 2.4 ) Evaluating effects of isolation on genetic and song distances using a landscape ecology framework

We evaluated the effects of different isolation modes (IBD, Isolation by Environment, IBE, and Isolation by Resistance, IBR) on genetic and song variation independently within a landscape ecology framework. For this, we used Multiple Matrix Regression with Randomisation (MMRR) (Wang, 2013), with significance assessed using 999 permutations (algatr package - Chambers et al., 2023). Location-level pairwise geographic distances (Isolation by Distance, IBD) were calculated as Haversine distances (geoSphere package; Hijmans et al., 2015) using sampling coordinates for both the song (7 locations) and genetic (13 locations) datasets. Landscape resistance (IBR) was based on the inverse of a habitat suitability model generated for S. albiventris (following Arunima et al., 2026, Figure 1d) and converted to pairwise resistance distances using the least-cost path function (gdistance package; van Etten, 2017). When both IBD and IBR were significant, resistance distances were regressed on geographic distance (lmer package), and the residuals were used to account for spatial autocorrelation, since geographic distance and resistance distances were highly correlated (r = 0.88). As the landscape exhibits an east-west precipitation gradient (Figure 1e), the environmental distance matrices (IBE) were derived from eight precipitation-related Bioclim variables (CHELSA v2; Bio12–Bio19). Principal component analysis was performed, and the first three principal components, which explained>90% of the variation, were retained (Supp. Table 3).

We tested eight response matrices (genetic: FST, Dps; song: 1-, 2-, and 3-gram presence-absence and weighted abundance) against three predictors (IBD, IBE, IBR), yielding 24 independent MMRR models (section 2.4). Additionally, we conducted partial Mantel tests (vegan Oksanen et al., 2024) to assess the relationship between song and genetic distances while controlling for geographic distance.

## Results

We used song data (87,589 notes) from 90 individuals and genetic data (10,638 SNPs) from 207 individuals to examine patterns of variation across a heterogeneous landscape.

### 3.1.1 ) No variation in spectral variables, but variation in syntax and ordering is evident

There was no distinct clustering of songs across locations when considering frequency-time parameters and the proportion of Complex Vocal Mechanisms together (Supp. Figure 3, PCA loadings: Supp. Table 4). Variance partitioning of frequency and time song-level variables revealed that within-individual variation accounted for the largest proportion of variance (83.18%), followed by between-individual variation (15.28%). Variance attributable to the population level was negligible, 1.54% (Supp. Table 5). When considered separately, the proportion of Complex Vocal Mechanisms in song syntax did vary significantly across locations, showing a decrease from west (GR) to east (KD)(GLMM beta distribution: β = -0.089, z = -2.90, p = 0.004), with individuals explaining ∼70% of the variation and population explaining ∼30% of the variation) (Figure 2).

**Figure 2:**
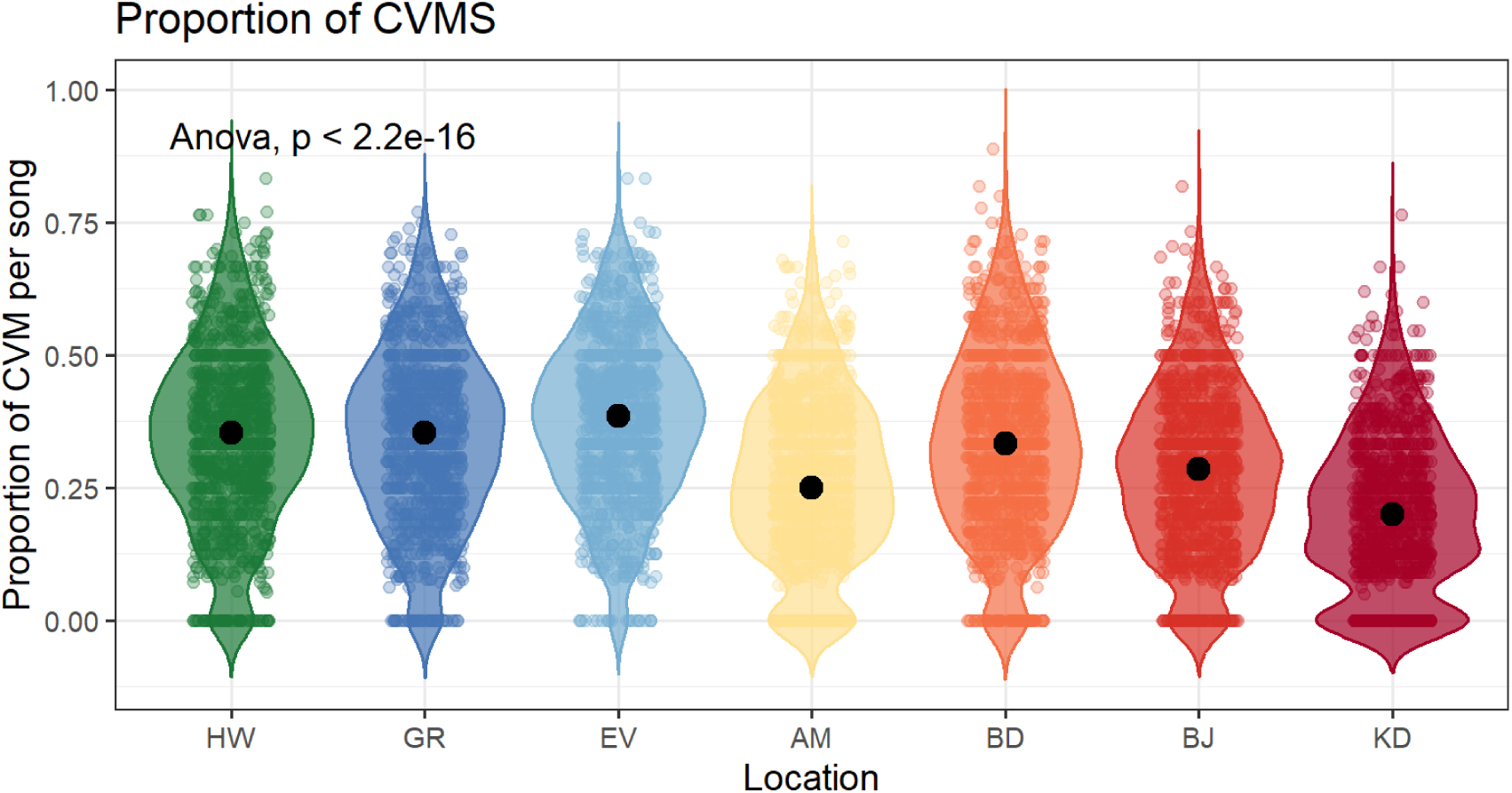
Showing the proportion of CVMs (Complex Vocal Mechanisms) in songs across locations, showing a higher proportion of CVMs in the western locations (GR and EV) as compared to the eastern populations (BJ and KD). The southernmost population HW also shows a high proportion of Complex Vocal Mechanisms.

**Figure 3:**
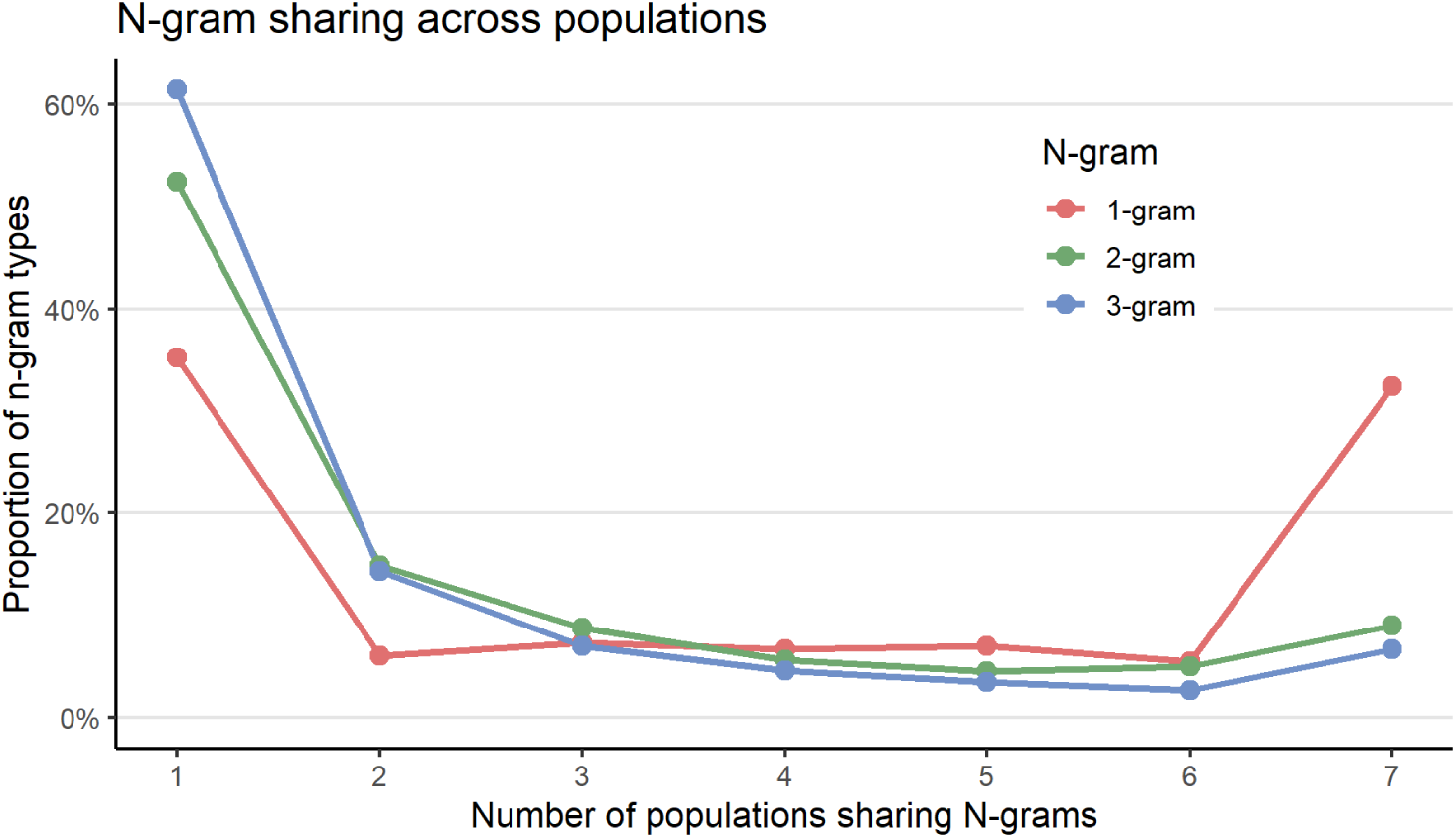
N-gram sharing across populations, ranging from N-grams unique to a single population (number of populations sharing = 1) to N-grams present in all populations (= 7), with intermediate values (2–6) reflecting partial sharing across a subset of populations. Basic note types (1-grams, red) show a bimodal distribution, with 35% unique to a single population and 33% shared across all seven. By contrast, 2-grams (green) and 3-grams (blue) show a much higher proportion of population-unique types and lower sharing across all populations, indicating that multi-note combinations are more population-specific than individual note types.

After notes were classified and filtered, there were 315 distinct notes (1-grams) (32% shared across all seven populations and 35.2% were restricted to a single population). Thus, the 1-gram distribution was bimodal, with notes either common across all populations or restricted to a single population (Figure 3, Supp. Figure 4a). Most 1-grams present across all populations were identified as flat introductory whistles that occur at the beginning of each song (Figure 4). On the other hand, longer combinations of notes (2-grams and 3-grams) were largely restricted to one population (2-gram 52.4 %, 3 grams 61.4%) while sharing between multiple locations (2-7) greatly decreased (proportion of N-grams shared between 7 populations: 2-grams 8.9% and 3-grams 6.15%, Figure 3, Supp. Figure 4b-c).

**Figure 4:**
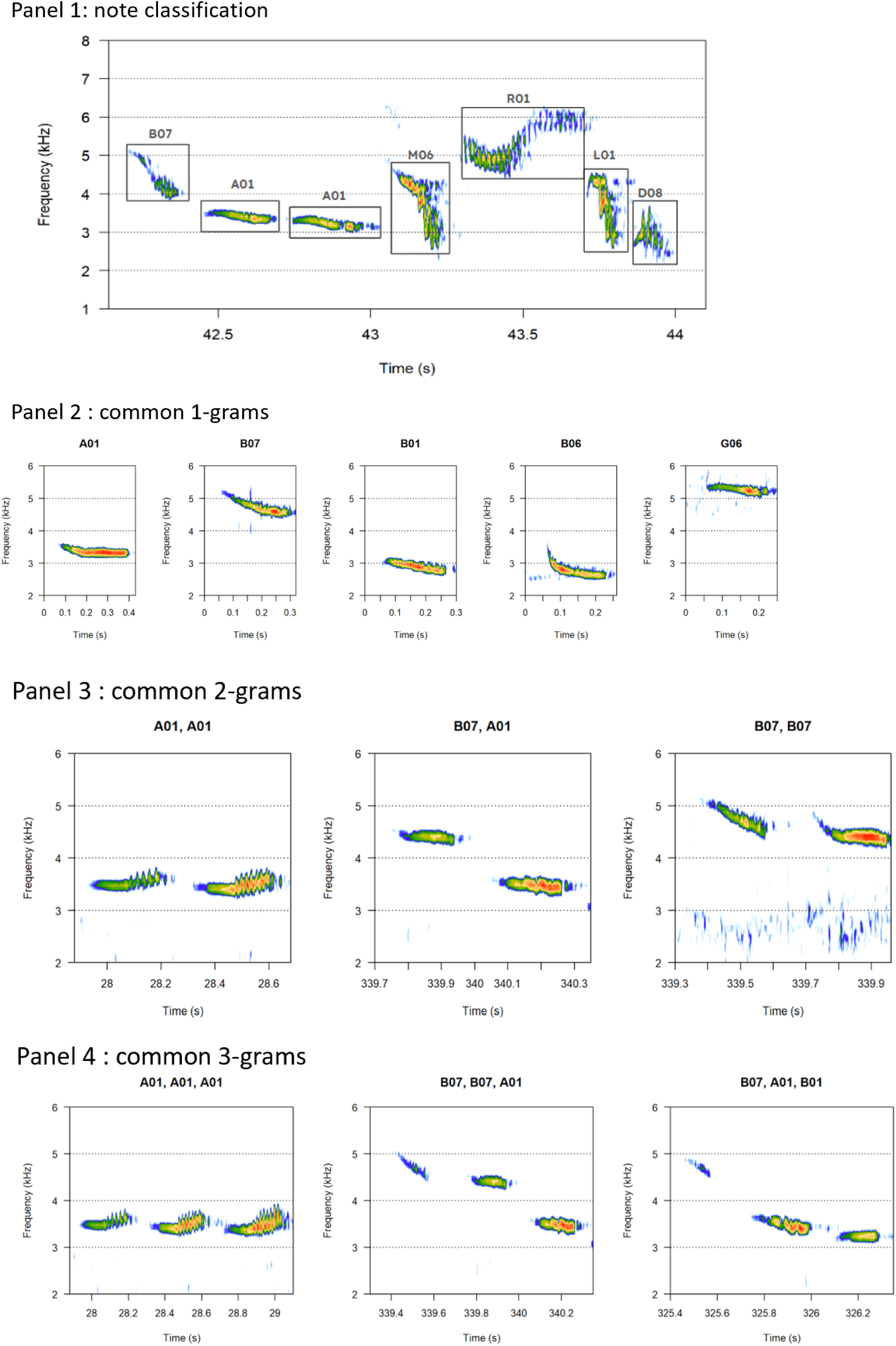
The first panel shows a song with N-gram classification, panel 2 contains the top 1-grams, panel 3 has the top 2-grams, and panel 4 has the top 3-grams.

We found that the observed n-gram sharing patterns depart significantly from a random combinatorial null model, keeping note frequencies and song lengths of an individual constant while randomly shuffling note order within a song (Supp. Figure 5). Two lines of evidence indicate that note combinations are non-randomly concentrated into a shared core repertoire rather than diverging across populations. First, sequences unique to a single population were rarer than expected by chance at both the 2-gram and 3-gram level (2-grams: observed 52.4% vs. null 61.7%, Z = −27.5, p < 0.001; 3-grams: observed 61.5% vs. null 82.6%, Z = −131.6, p < 0.001), indicating less population-specific syntax than random combination would generate. Second, sequences shared across all seven populations were correspondingly more common than expected (2-grams: observed 8.9% vs. null 4.2%, Z = +53.2, p < 0.001; 3-grams: observed 6.7% vs. null 1.0%, Z = +184.9, p < 0.001), with universal sharing roughly two-fold and seven-fold above the null expectation at the 2- and 3-gram level, respectively. Together, these results show that S. albiventris song syntax is organised around a set of universally shared note combinations, a structure not explained by note-level sharing alone.

**Figure 5.**
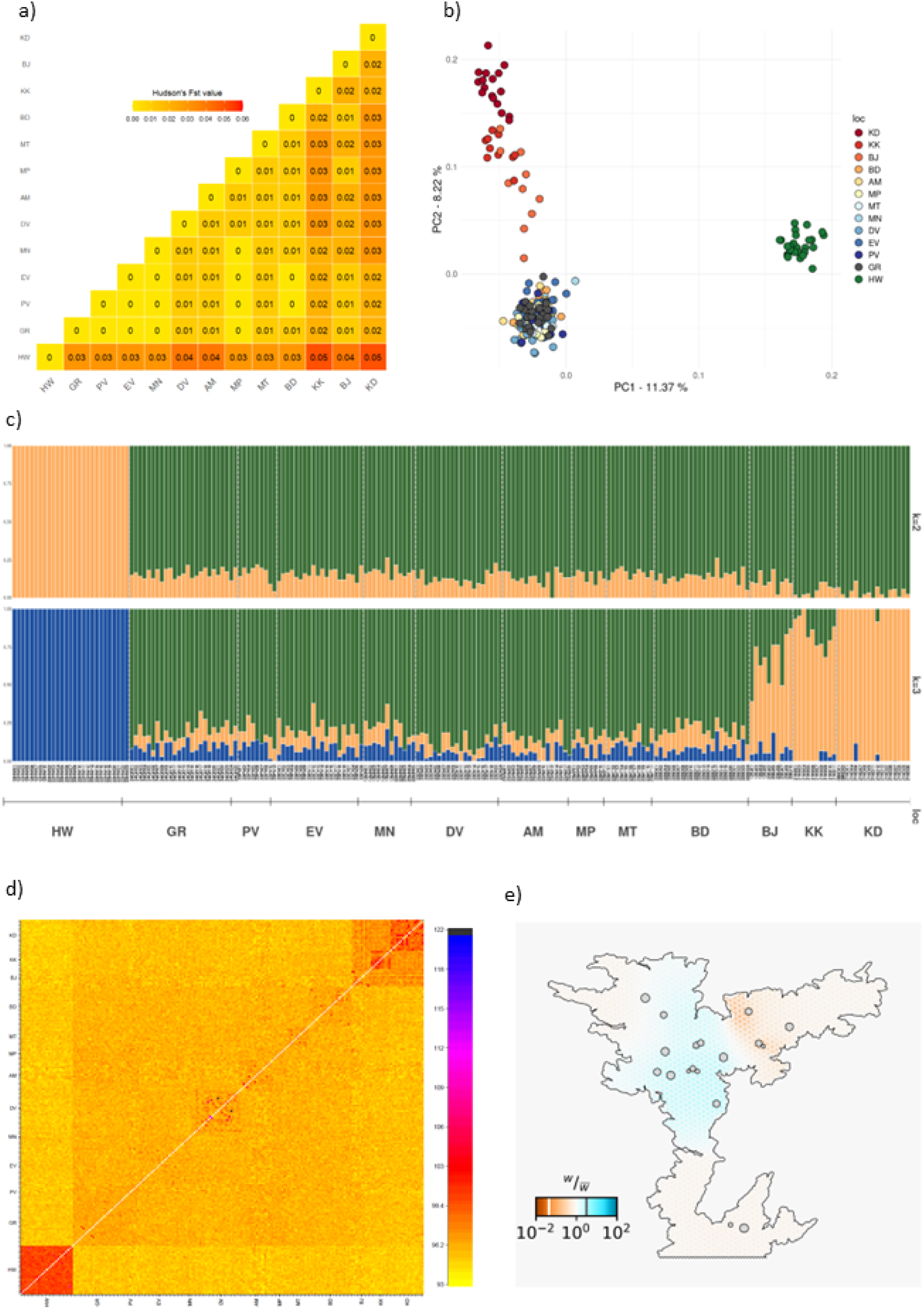
a) Pairwise FST matrix for all populations sampled, b) Principal Component Analysis and Discriminant Analysis of Principal Components for genomic data, c) Structure plot of shared alleles for K =2 and K = 3, d) Co-ancestry plot for shared haplotypes using fineRADstructure e) FEEMS plot showing estimated effective migratory surfaces highlighting greater deviation from IBD showing higher than expected effective migration (blue) and lower than expected effective migration (red) indicating presence of some barriers.

### 3.1.2 ) Populations differ across all levels of N-gram sharing; no effect of geographic distance on song sharing

We found weak but significant differences in local-level syntax (whether scored as presence-absence or abundances of grams) across all N-gram levels. The differences between locations declined from 1-gram to 3-gram for presence absences *(PERMANOVA presence-absence: 1-gram R² = 0.227, F = 4.056; 2-gram R² = 0.157, F = 2.569; 3-gram R² = 0.157, F = 2.575; all p < 0.001)* and abundances (PERMANOVA abundances 1-gram R² = 0.165, F = 2.732; 2-gram R² = 0.150, F = 2.448; 3-gram R² = 0.148, F = 2.407; all p < 0.001). These differences persisted even after correcting for potential sample-size bias (effect sizes in Supp Table 6).

Within-population individual variability in syllable repertoires was significantly heterogeneous across populations at 1-gram and 2-gram levels for both presence-absence and abundance matrices (PERMDISP: presence-absence: 1-gram p < 0.01, 2-gram p = 0.004; abundance: 1-gram p = 0.045, 2-gram p = 0.021; Supp. Figure 6), indicating that some populations contain more individually distinct singers than others. This heterogeneous dispersion means that observed between-population differences at these levels (see PERMANOVA above) may partly reflect unequal within-population individual variability rather than the observed differences between population centroids (PERMANOVA results). At the 3-gram level, individuals within locations did not differ in dispersion (PERMDISP: presence-absence: p = 0.016; abundance: p = 0.054), indicating differences are entirely due to differences in locations (PERMANOVA results above). Individual compositional distinctiveness (distance of an individual to population centroid) increased with n-gram order (for presence absence and abundances), indicating that note sequence and transition structure carry more individually distinguishing information than note identity alone (presence absence beta GLMM; 2-gram vs. 1-gram: β = 0.796 ± 0.021, z = 38.35, p < 0.001; 3-gram vs. 1-gram: β = 0.907 ± 0.021, z = 43.67, p < 0.001; abundances beta GLMM: 2-gram vs. 1-gram: β = 0.680 ± 0.049, z = 45.65, p < 0.001; 3-gram vs. 1-gram: β = 0.94201 ± 0.0.014, z = 63.64, p < 0.001). Variance partitioning indicated that this distinctiveness was driven predominantly by individual rather than population-level differences (presence-absence individual-level variance = 0.041 vs population-level variance = 0.0065; abundances individual-level variance = 0.05 vs population-level variance = 0.004).

**Figure 6:**
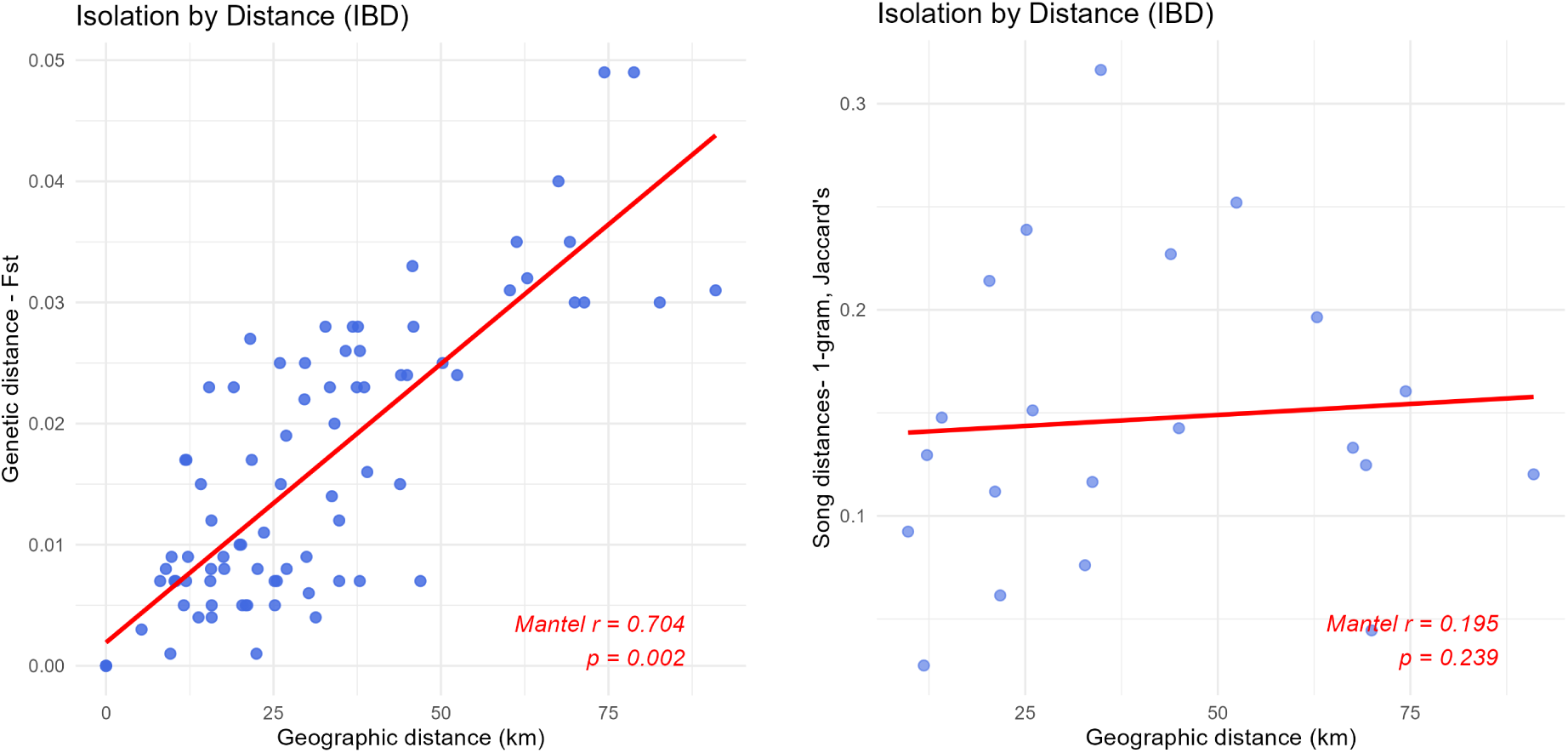
Relationship between geographic distances and genetic distances (a) and song distances (b). Genetic distances show a strong isolation-by-distance pattern, whereas song distances (here using a 1-gram presence-absence matrix) are not.

### 3.2 ) Playback experiments - birds do not discriminate between local and non-local songs

We observe that frequency-time parameters in songs across sites are not significantly different, whereas locations show differences in syntax. To test discrimination towards local and non-local songs, we conducted a playback experiment in one location - Kodaikanal, across two seasons.

We found that behavioural responses to stimuli segregated into two categories: vocal responses (PC1, 43.08% of variance explained) involving latency to sing and song duration, and physical responses (PCA: PC2, 26.67% of variance explained) including flight frequency and distance (Supp Table 7). We find that seasonal variation did not influence behavioural responses to playback stimuli. Neither vocal (PC1) nor physical (PC2) responses differed between breeding and non-breeding seasons (PC1: t-test, Bonferroni corrected t = 0.87, df = 132.43: p = 0.39; PC2: t-test, bonferroni corrected = t = -0.83, df = 131.43, p = 0.41). Considering this result, the season was excluded from subsequent models as a variable.

Testing for behavioural differences in responses to local and non-local (near and far) stimuli, we find that birds showed the strongest vocal responses to local stimuli (reference category; intercept = 0.20 ± 0.27 SE). Responses were significantly reduced toward control stimuli compared to local stimuli (LMM: β = –1.67 ± 0.30 SE, t = –5.59, p < 0.0001). Responses to non-local near and non-local far stimuli were not significantly lower than responses towards local stimuli (non-local near: β = –0.38 ± 0.22 SE, p = 0.080; non-local far: β = –0.35 ± 0.22 SE, p = 0.118; Table 1). Individual identity accounts for a very large proportion of variance in vocal responses, i.e., birds respond differently to any given stimulus irrespective of which location the stimulus is from (Conditional R2 = 0.546, marginal R2 = 0.122). Physical responses (PC2) also did not differ significantly across playback stimulus types. Physical responses to non-local near, non-local far, and control stimuli were not significantly different from local stimuli (non-local near: β = –0.18 ± 0.22 SE, p = 0.430; non-local far: β = –0.28 ± 0.23 SE, p = 0.220; control: β = –0.36 ± 0.31 SE, p = 0.245; Table 2). Unlike the vocal response model (PC1), the physical effect model (PC2) shows that the random effects contribute relatively little additional explanatory power, suggesting that neither stimulus type nor individual accounts for much of the variation in physical responses (Conditional R2 = 0.077, marginal R2 = 0.014). When vocal and physical response variables were examined independently, none differed significantly across local, non-local near, and non-local far stimuli (ANOVA singing duration: F₂,₁₁₈ = 0.695, p = 0.211; latency to sing: F₂,₁₁₈ = 0.871, p = 0.421; average distance to speaker: F₂,₁₁₈ = 0.537, p = 0.586; flights over speaker: F₂,₁₁₈ = 1.406, p = 0.249).

### 3.3 ) Low genetic differentiation is seen between populations, though population structure is present and is consistent with isolation by distance

Genetic data processing resulted in 207 of the 242 individuals from the 13 locations for genetic population structure analysis. After filtering, we retained 10,638 SNPs. We found the Highwavies (HW) population in the south was the most genetically distant from the other sampled locations in the Anamalai-Palani, with the highest differentiation observed between HW and Kodaikanal (KD) and Kookal (KK) in the east (pairwise Hudson’s FST of 0.049; Figure 5a). We find patterns of genetic structure highly concordant across PCA and ADMIXTURE. HW is seen segregating separately from the other sampled sites in the Anamalai-Palanis (PCA results - Figure 5b) as well as in the admixture plot (Figure 5c, ADMIXTURE, K = 2, 3, and clustering analysis). However, this differentiation is minor, and the populations are mainly mixed (as K = 1 yields the lowest cross-validation error score in ADMIXTURE).

Co-ancestry results (fineRADstructure) also indicate that individuals from the HW population had greater haplotype sharing than individuals from the other populations (Figure 5d). Estimating migratory surfaces (Figure 5e) also highlights locations on the western side of the Anamalai-Palai hills with higher-than-expected effective migration, while the eastern locations of BJ, KK, and KD show lower estimated effective migration (in brown). The southernmost location of HW (in white, Figure 5d) shows a near-IBD result with no evidence of increased or decreased migration.

### 3.4 ) Song variation is governed by IBD, IBE, and IBR, though historical genetic differentiation strongly follows IBD

We tested for the effects of isolation on genetic variation and song differences. Pairwise genetic differentiation, calculated as Fst, across 13 sampled locations showed a strong and significant relationship with geographic distance (Isolation by Distance; IBD) (Supp Table 8a, Figure 6a). The IBD model explained 61% of the variation in genetic distance (R² = 0.61; F = 118.16; p < 0.001), with a significant positive effect of geographic distance (β = 0.78, 95% CI: 0.66–0.90). Even after excluding the furthestmost population in the south (HW), there is a strong positive association of geographic distances and genetic distances (β = 0.58, 95% CI: 0.39–0.74, R² = 0.32; F = 30.05; p < 0.001). This indicates that genetic differentiation increases substantially with increasing spatial separation. In contrast, environmental variation explained via precipitation differences did not significantly explain genetic structure. The precipitation model (IBE) explained only 7% of the variance (R² = 0.07; F = 1.98; p = 0.54), and none of the predictors was statistically significant. Isolation by Resistance (IBR) was also significantly associated with genetic differentiation when considered independently (R² = 0.50; F = 75.59; p < 0.001), with a strong positive effect of resistance distance (β = 0.71, 95% CI: 0.57–0.84). However, in a combined model of geographic (IBD) and residual resistance distances (IBR) model, we find resistance does not explain additional genetic variation beyond that captured by geographic distance alone (R² = 0.62; IBD: β = 0.78, p < 0.001; IBR: β = 0.12, p = 0.40). Overall, these results indicate that geographic distance is the primary driver of genetic structure in the study system, with little support for effects of environmental variation or landscape resistance. When evaluating genetic distances in terms of DPS (i.e., contemporary genetic distances), we found that none of the isolation-by-geographic-distance, environment, or resistance factors significantly predicted D_PS_ distances (Supp Table 8b).

In the case of songs, we find that neither geographic distance (Figure 6b), environmental distances, nor resistance distances explain variation in notes (1-gram) and 2-grams for presence-absence(Supp Table 8c) and abundance distances (Supp Table 8d). For 3-grams, geographic and resistance distances did not significantly explain variation in either presence-absence (Supp Table 8c) or abundance measures (Supp Table 8d). On the other hand, environmental distances were able to explain some variation in 3-gram presence-absence (PC1: β = 0.68, 95% CI = 0.39-0.98, p value = 0.01, R² = 0.55, p = 0.01, Table 8d), but not 3-gram abundance distances (Table 8c). Additionally, a negative, but non-significant, relationship between song and genetic distances was observed (partial Mantel test with FST and DPS), suggesting limited support for song variation driven by genetic differences (Supp. Table 9).

## Discussion

Our study, conducted across multiple populations using fine-scale song and genetic data, reveals that patterns of song sharing and genetic similarity are discordant.

### Combinatorial syntax highlights high individual-level variation and low population-level segregation

Birdsong variation can be complex to assess, particularly in open-ended learners, where it can be challenging to find an appropriate evaluation measure. In S. albiventris, songs from different populations are aurally indistinguishable and have similar frequency-time parameters. Yet syntactic analysis reveals structured patterns in note and note-combination sharing.

Certain notes (1grams) are common to all individuals (Price et al., 2008; Benedict and Bowie, 2009), possibly reflecting conformist bias driven by social influence (Lachlan et al., 2018). Longer note combinations are less shared, similar to syntax sharing in Puget Sound white-crowned sparrows (*Zonotrichia leucophrys pugetensis*) and nightingales (*Luscinia megarhynchos*) (Nelson et al., 2004) (Kipper et al., 2004);(Grießmann & Naguib, 2002), where the universal common elements coexist with population- and individual-level differences associated with higher-order syntax( Lee et al., 2019). Population-level differences account for only a small proportion of song syntax variation, with the majority of variation explained by individual-level syntax variation. In line with this, Sawant et al. (2025) monitored a few individuals of the white-bellied sholakili over three consecutive years and showed syntactic change, with syllables dropped and new ones incorporated. Similarly, within-individual variation greatly exceeds population-level differences in the Blue-throated flycatcher Cyornis rubeculoides complex when assessed using spectro-temporal parameters(Singh et al., 2020).

Weak population-level differences, tending toward the absence of dialects, align with patterns in other open-ended learners, such as nightingales (Jäckel et al., 2022). This may partly reflect the absence of prolonged geographic isolation, consistent with the isolation-by-distance pattern in population genetic structure. A high cultural rate of change may further mask dialect formation, contrasting with the pied flycatcher (Ficedula hypoleuca), an open-ended learner in which dialects are nonetheless present. Longitudinal individual-level recordings across locations would help clarify the stability of population- and individual-level signatures. However, longer, continuous stretches of individual-level recordings across locations, collected longitudinally, would be important in the future to gauge the stability/presence of population- or individual-level signatures.

Based on current evidence, we suggest that 1-grams shared across all individuals may have a stochastic basis rooted in social conformity, while higher-order syntax (2-grams and 3-grams) may reflect more deterministic, individual-level processes. Understanding the mechanisms underlying song variation in a tropical, sedentary, year-round territorial open-ended learner adds to a literature dominated by temperate and migratory species.

### Playback studies confirm no differential discrimination towards local and non-local songs

Our playback experiments revealed that birds did not discriminate significantly between local and non-local songs, suggesting a broadly similar behavioural response. This finding raises important questions about the function of song and recognition thresholds in species exhibiting low conformity. This result also validates the previous finding that, although syllables in songs across locations differ, the major driver is that individuals themselves have specific syllables. Given that songs across populations are spectrally similar, yet higher-order syntactic structures (i.e., 3-grams) confer individual-level signatures with limited sharing, it is possible that playback of non-neighbour songs—whether local or non-local—fails to elicit differential responses. Birds may respond only to neighbours, and we know that, within a location, syntax sharing decreases with increasing distance (within 1km) (Sawant et al., 2025). Another reason for non-discrimination may be that similar songs in frequency-time parameters, as well as 1-gram sharing, drive recognition and response. In olive sparrows (*Arremonops rufivirgatus*), for example, one population responded more strongly to local songs, whereas another showed equal responses to both local and non-local calls (Fernández-Gómez et al., 2012), while in pied flycatchers *(Ficedula hypoleuca)*, another bird with complex songs, individuals show stronger responses towards local songs as opposed to foreign songs(Gallego-Abenza et al., 2025). These contrasting outcomes emphasise the need to replicate playback experiments across different locations, as behavioural responses observed in one location may not generalise across a species’ distribution.

### Geographic distances drive patterns in genetic differentiation

The recovery of fine-scale genetic structure driven primarily by isolation-by-distance and a relationship with geographic distance is new in this system. This contradicts a previous study (Robin et al 2015), which also found high contemporary differentiation (D_PS_) in the landscape and different patterns of genetic population structure. Perhaps these differences are due to the microsatellite markers (previous study), which are multi-allelic, unlike SNPs (biallelic filtered; this study). Higher mutation rates may indicate lower allele sharing and greater differences between locations, as also suggested by Haasl and Payseur ((Haasl & Payseur, 2011). Such differences in population structure using microsatellite loci and SNPs have also been recovered in the Gunnison sage-grouse (*Centrocercus minimus*) (Zimmerman et al., 2020) and Crucian carp (*Carassius carassius*) (Jeffries et al., 2016), where SNP clustering analyses revealed strong demographic independence that was not recovered by microsatellite data.

We also find low effective migration estimated from genomic data across the eastern sampled locations (KD, KK, and BJ), which are consistent with recent connectivity inferred from occupancy data (Arunima et al., 2026). Similar patterns have been recovered in Amazonian birds where historical barriers persist as low-migration surfaces despite recent landscape dynamism (Berv et al., 2021; Musher et al., 2022). Genetic population structure was consistent across methods (Robin et al. 2015; present study), revealing isolation by distance across a mid-elevation 1000m ridge (connecting HW and Anamalai-Palani). A 2023 survey (by CA and SS) on the ridge detected no *S. albiventris* in the forest patches, consistent with genomic evidence indicating that this tropical montane endemic species is unsuitable for this habitat. Overall, this result highlights that across a small spatial scale, such as the sky island system in the tropics, topographical barriers drive genetic structuring in montane endemics, and recent anthropogenic change fails to blur the differences yet, partly because secondary contact and genetic exchange may take time to accumulate and cannot be captured with the current methods.

### Discordance in song and genetic population structure

Our findings reveal clear variation in both song and genetics; however, these patterns do not align. While genetic distances show a strong correlation with geographic separation, song variation does not at similar spatial scales, likely because cultural change is more dynamic than that of inherited genes. The discordance between cultural (song) and genetic variation aligns with research on birdsong and genetic variation (Poesel et al., 2017; Searfoss et al., 2020; Soha et al., 2004). Often, song variability depends on the measures used in evaluation, and in species with large repertoires, we find note type and note organisation to be important (Briefer et al., 2008; P. Singh & Price, 2015). In such species, although territorial and non-social, individuals exhibit song sharing with neighbours, sharing more units of syntax with neighbours than with non-neighbours (Sawant et al., 2025), a phenomenon also observed in skylarks, another species with a complex repertoire (Briefer et al., 2008). This may be to maintain territories, as we know that individuals are year-round territory holders. We had expected locations closer together, especially those connected by woodland expansion and increased connectivity (BJ, KK and KD) (Arunima et al., 2026), to share songs due to a decrease in resistance between locations, yet we have no such patterns. Song variation, especially at higher-order note strings (2-gram, 3-gram), becomes more individual-specific, with reduced sharing even within a population. Although habitat fragmentation is a common explanation for song-sharing patterns globally (Pérez-Granados, Osiejuk, and López-Iborra 2016; Pavlova et al. 2012), this study reveals that recent connectivity is not reflected in song- and genetic-sharing patterns. Song variation remains unexplained by geographic distance, environmental and habitat variation, which may be due to the complex nature of cultural traits (Williams, 2021), which may have a stronger social selection component compared to various factors of resistance tested here (IBE, IBR, or IBD). Social selection pressure may also drive the evolution of complex communication signals (Lachlan & Servedio, 2004; Rose et al., 2018).

## Conclusion

Our study suggests that there is no linear association between cultural variation and genetic connectivity in a species with high cultural variation. In an open-ended Passeriform like S. albiventris, the song repertoire comprises a core, global set of basic units and note combinatorics, resulting in weak population signatures but strong, unique individual signatures. We observe that geographic distance drives genetic differentiation; however, song sharing is not explained by isolation-by-distance or by environmental and habitat resistance. The high variation in song may be due to cultural drift or learning constraints, with even a component of individual recognition or sexual selection, which are avenues to be tested.

## Acknowledgements

We thank the Kerala and Tamil Nadu State Forest Departments for animal capture permits and forest accommodation. For fieldstation and logistics support we thank Kodaikanal International School and Hume Centre for Ecology and Wildlife Biology. This work would not have been possible without the field team and we thank - Kamaraj, Subhash M, Aparna Rao, Vinay K L, Shivam Shinde, Yuvraj, Meghana RJ, Viral Joshi, Mubeen M, Archita Sharma, Aditya Panigrahy, Amrutha Rajan, Helly Vyas and Sethulakshmi Nair. We thank Akashram for helping us optimise the n-gram classification script. We thank Arunima Jain and Akshay Herur for generating the resistance layer and highest precipitation quarter layer for analysis and figures. CA would also like to thank Vinay K L, Jobin Varughese and Meghana Natesh for discussions during the development of the project and for feedback. CA would also like to thank members of her thesis advisory committee for discussions that helped fine-tune aspects of this study.

---

**Supp Fig 1:**
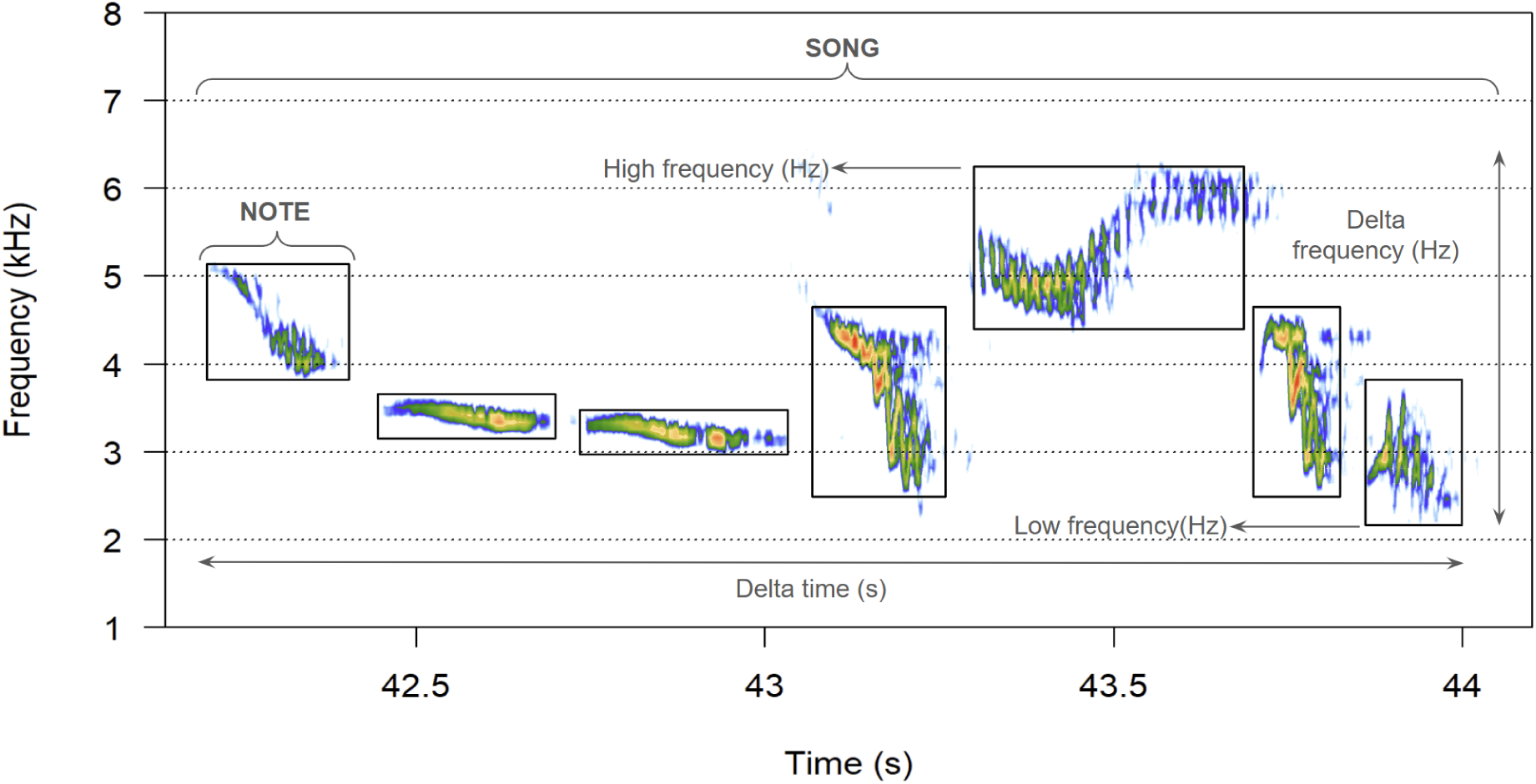
A spectrogram of a song of S. albiventris, highlighting the frequency and time parameters.

**Supp Figure 2:**
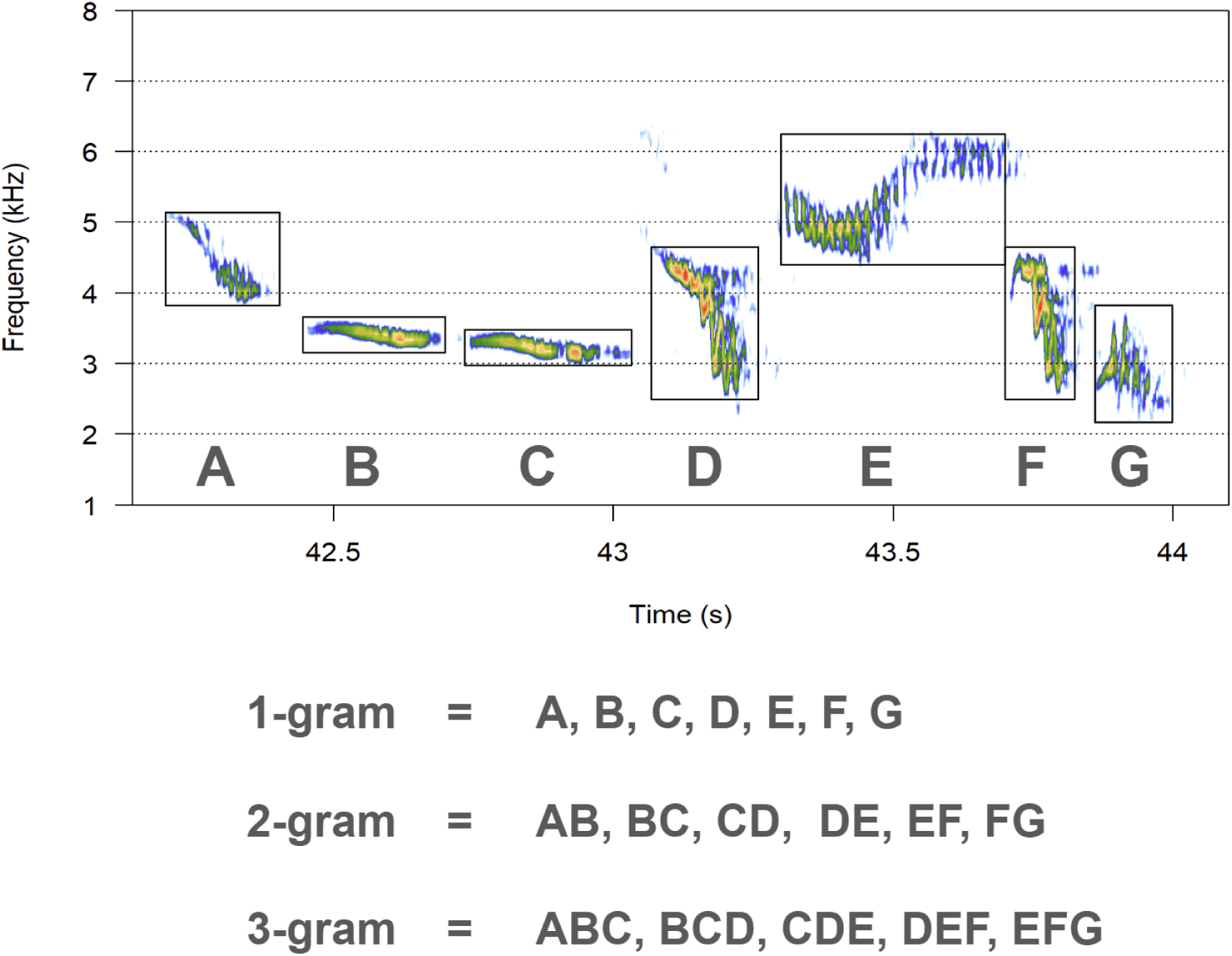
N-gram extraction at the 1-gram, 2-gram and 3-gram level for a song in the white-bellied sholakili

**Supp. Figure 3:**
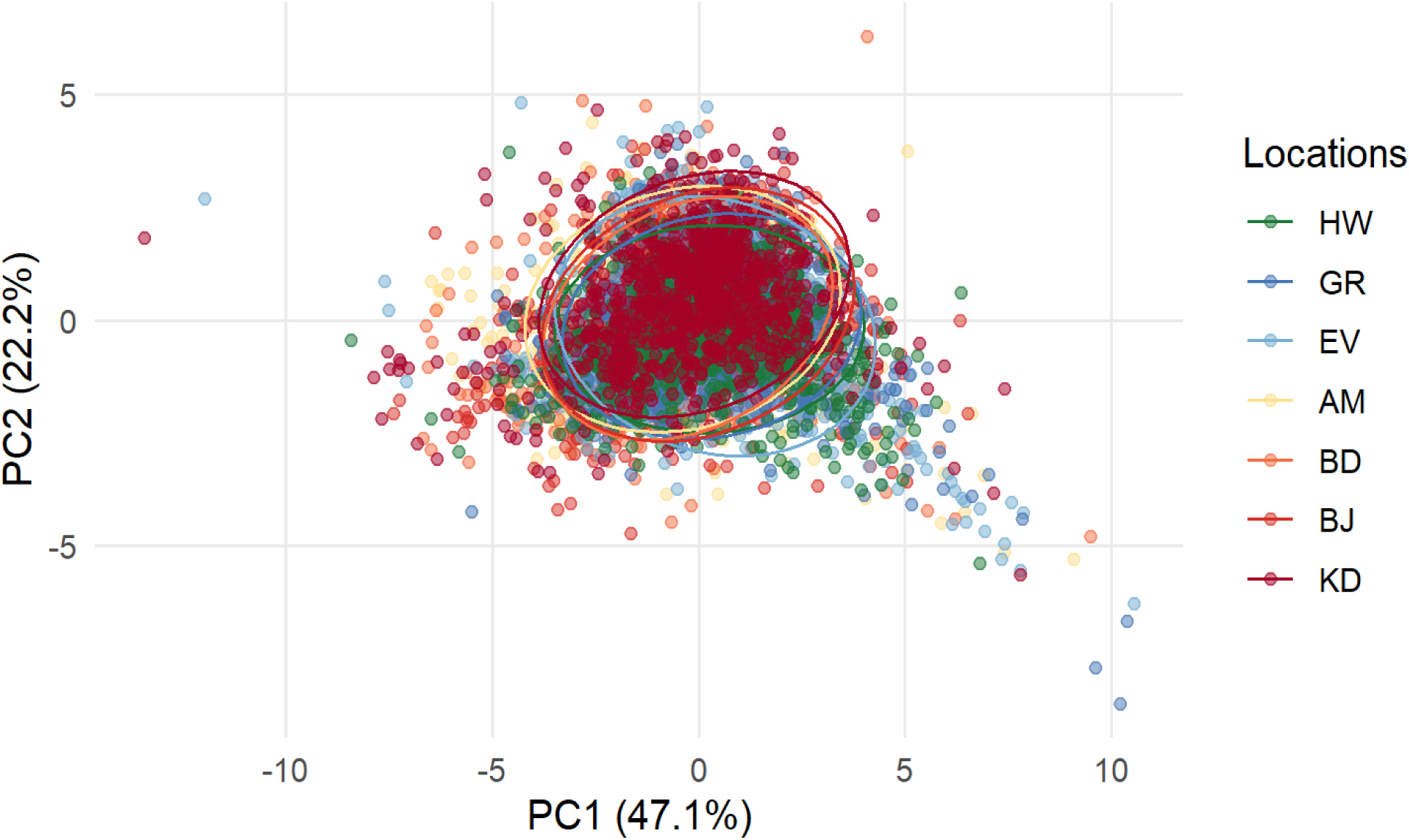
PCA score plot of song level eight acoustic parameters, including spectral, temporal, and the proportion of Complex Vocal Mechanisms (CVMs) across seven sampled populations, with each point representing one song colored by population of origin.

**Supp. Figure 4:**
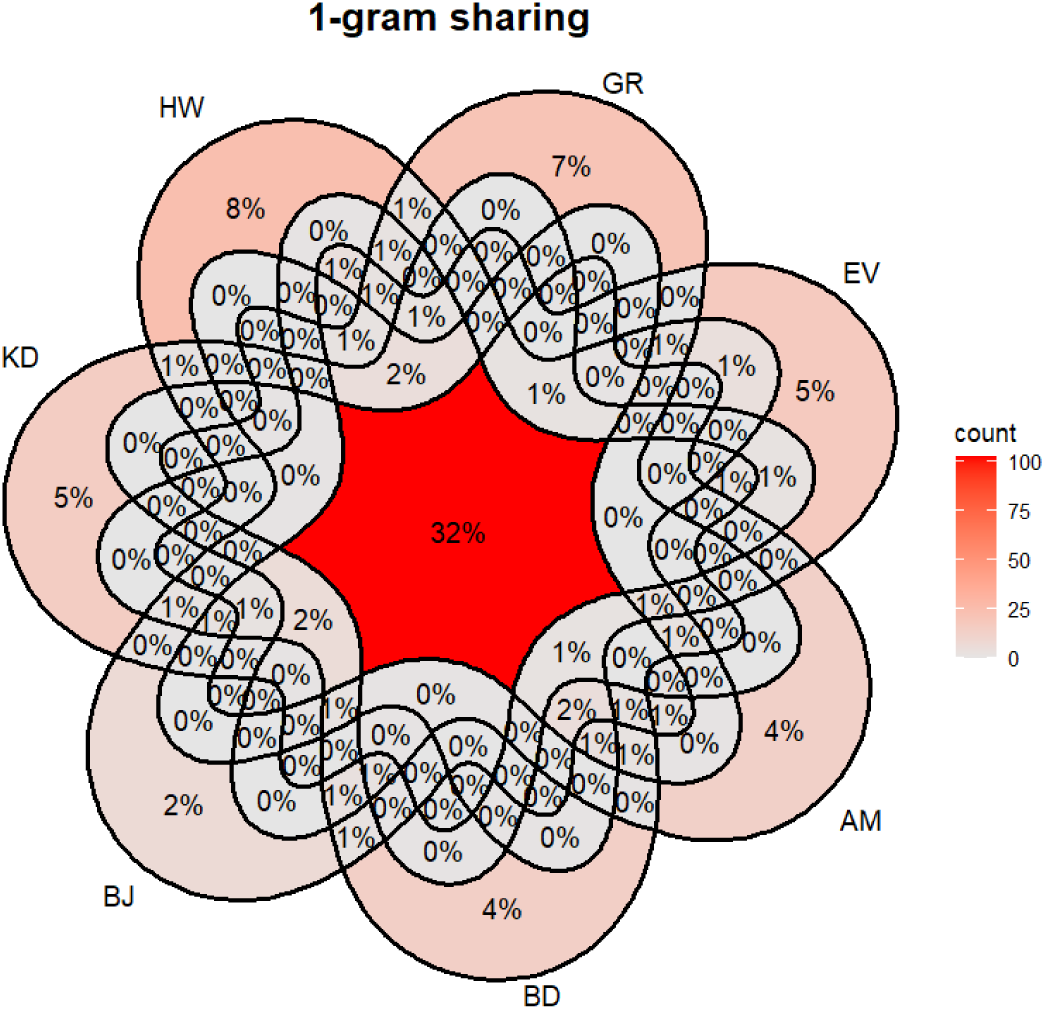
a)Venn diagram showing 1-gram sharing across the seven sampled population *b)* Venn diagram showing 2-gram sharing across the seven sampled populations *c)* Venn diagram showing 3-gram sharing across the seven sampled populations

**Supp. Figure 5:**
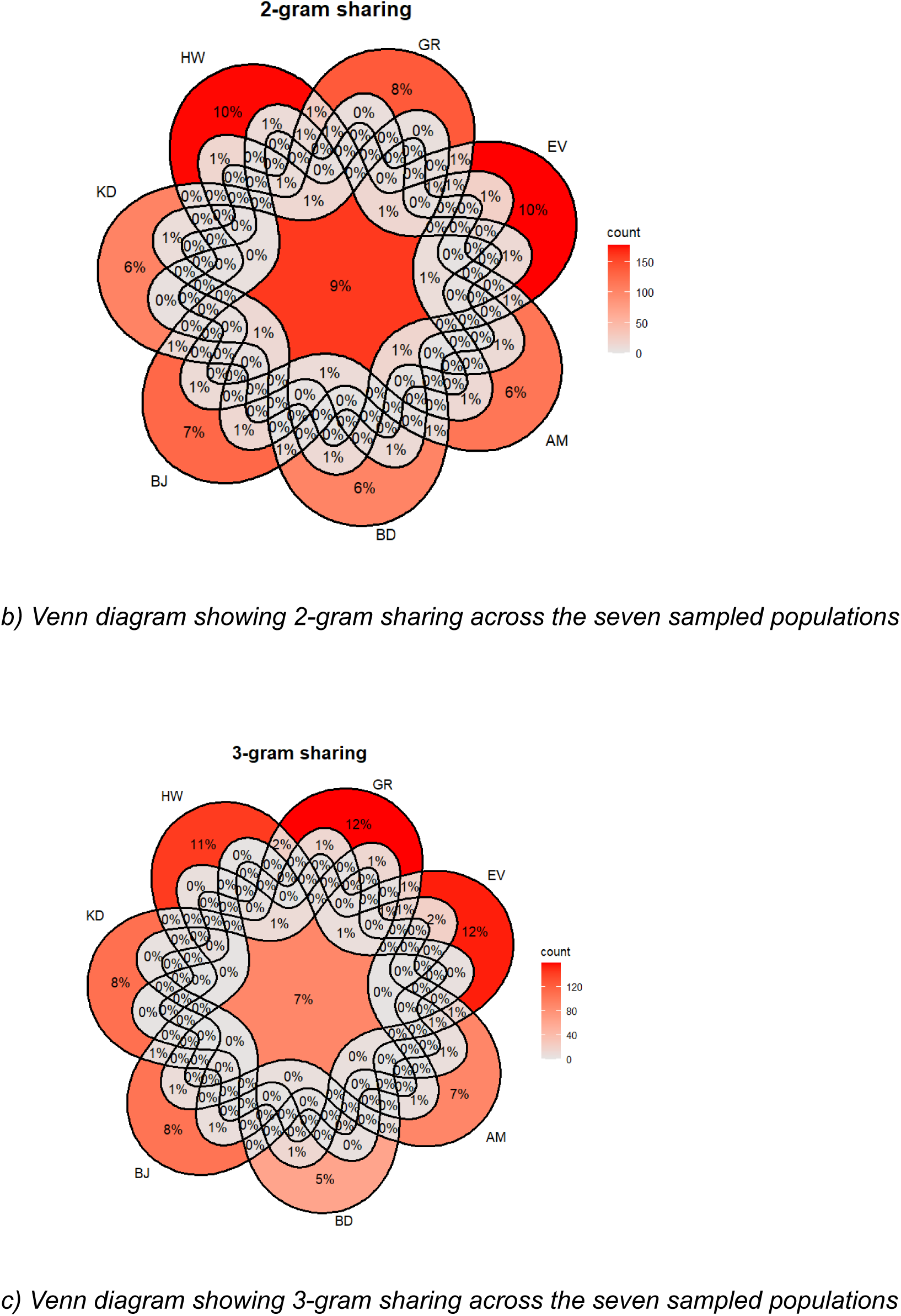
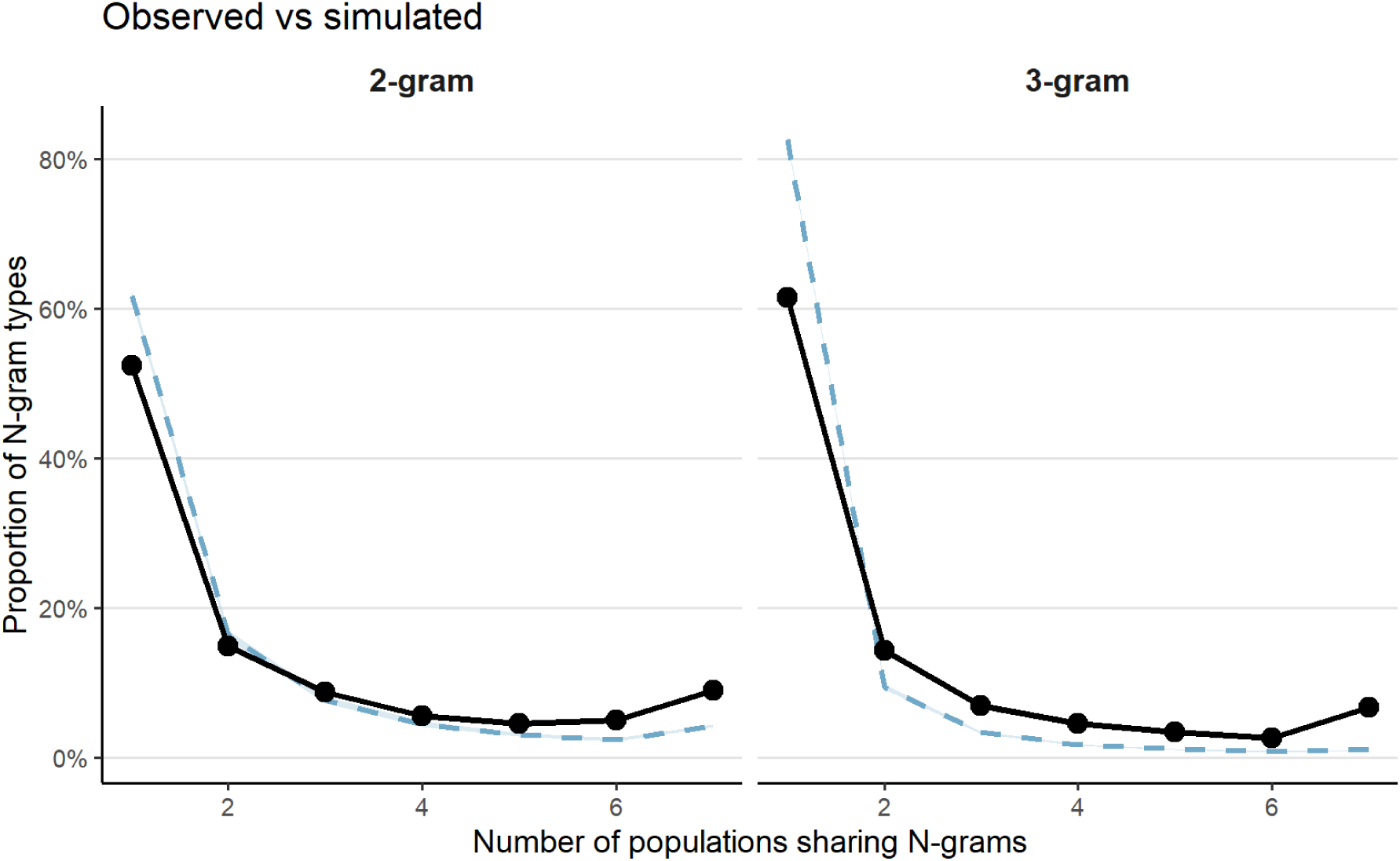
Observed (black) and simulated (blue dashed) proportions of 2-gram (left) and 3-gram (right) sharing across populations. Simulated n-grams were generated by preserving the note pools and song number for an individual and randomly shuffling the notes to generate N-grams. Proportions of 2-gram and 3-gram unique to a population were significantly lower than simulated values, while universal 2-grams and 3-grams (i.e., shared across all populations) are greater than simulated combinations.

**Supp Figure 6.**
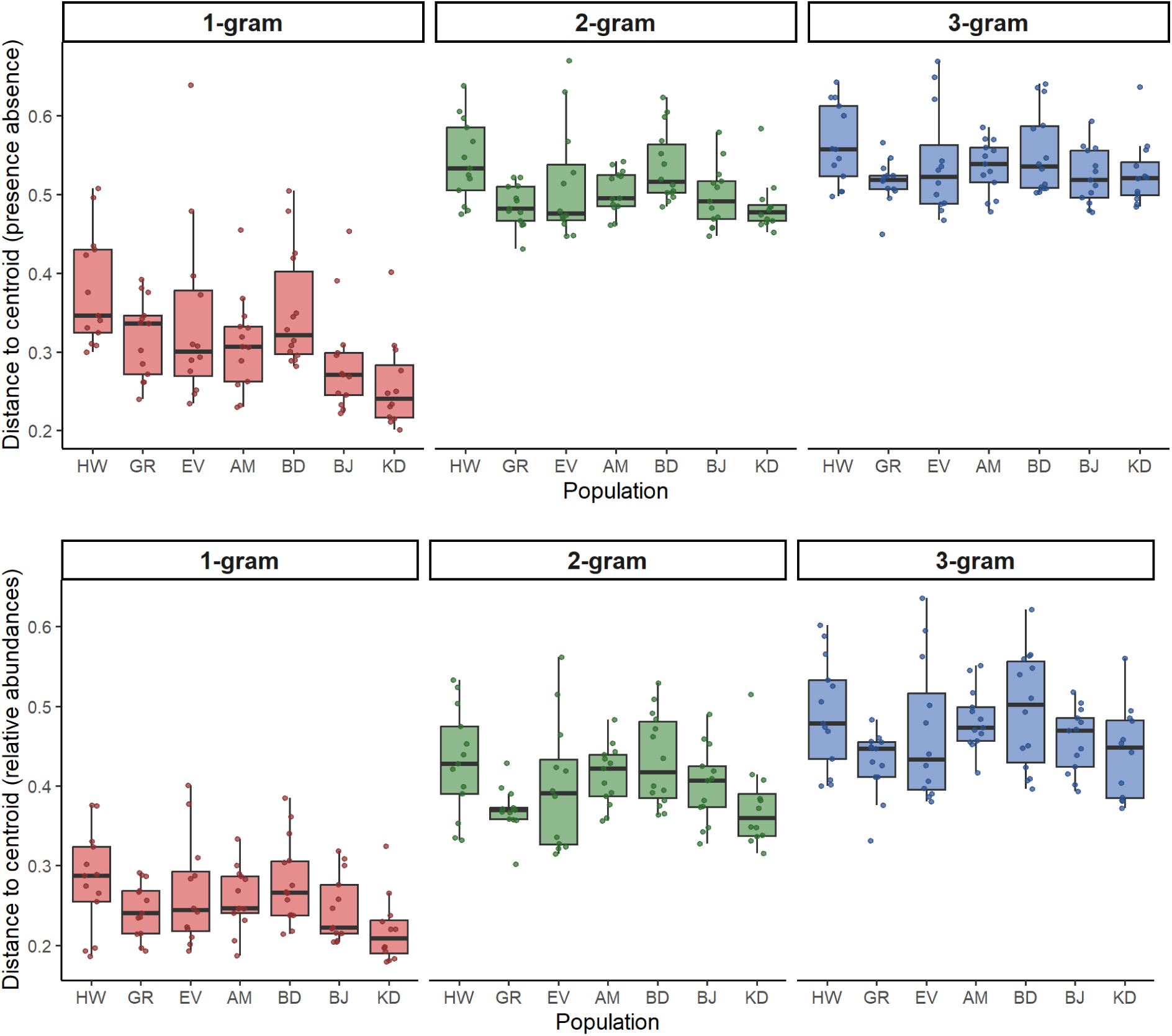
Box plots indicate dispersion distances of an individual (dots) to the population centroid (0) across N-grams (1-gram to 3-grams) with presence-absence distances (top panel) and relative abundances (bottom panel). Dispersion distances of individuals from their location centroids increase with increasing N-gram order.

**Supp Figure 7:**
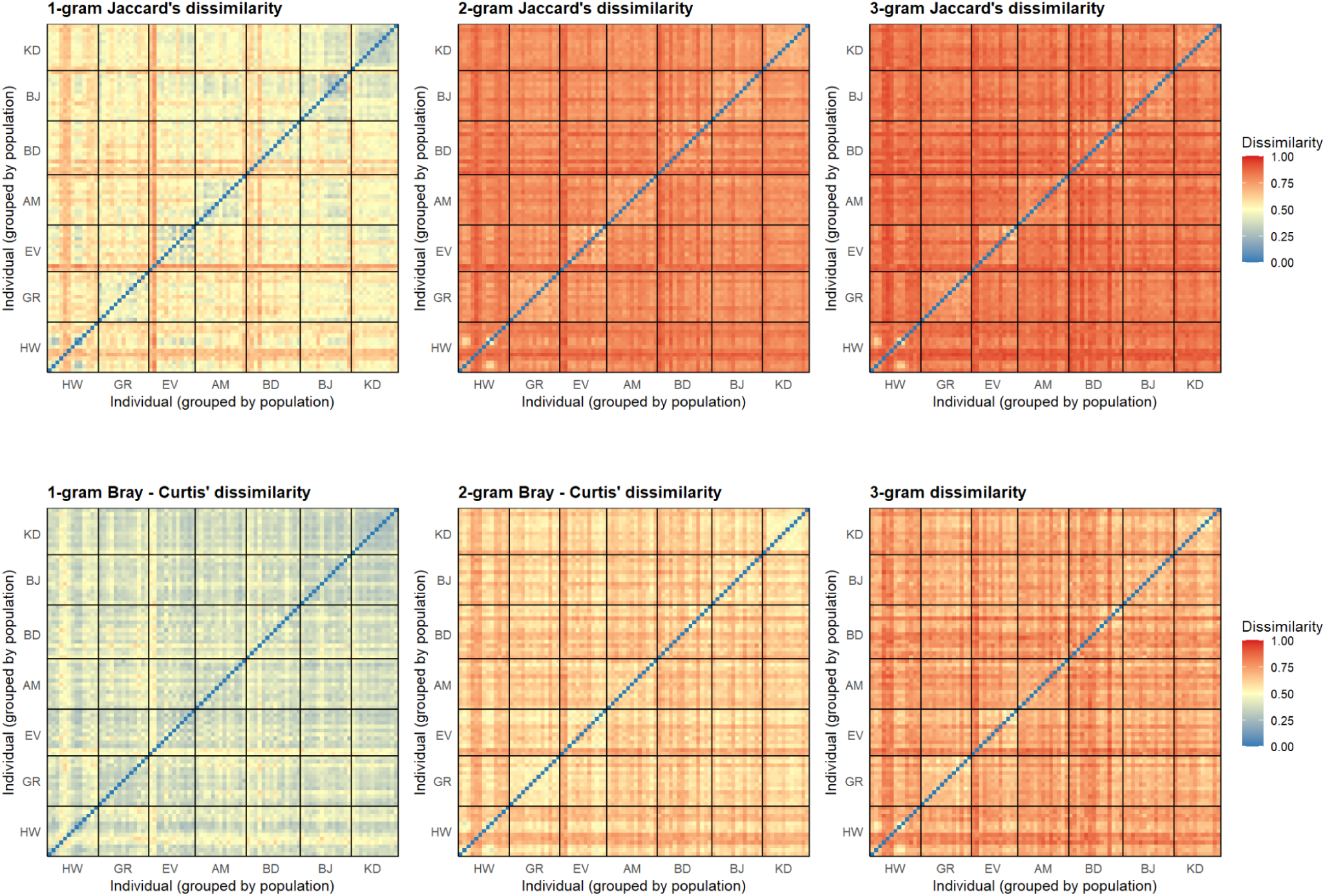
Heatmaps highlighting individual-level pairwise dissimilarity of N-gram sharing using presence -absence (Jaccard’s distance) and relative abundances (Bray-Curtis distance) of notes. The trend across both distances shows that dissimilarity between individuals increases as combinatorial N-gram order increases.

**Supp. Table 1:** Summary of all captured blood samples and ddRAD sequenced individuals, old (2005-2012) and new (2018-2024). 332 individuals were captured, of which 242 were sequenced.

|  | Captured | Sequenced | After filtering for missingness<br>& relatedness |
| --- | --- | --- | --- |
| **AM** | 21 | 21 | 16 |
| **BD** | 52 | 23 | 22 |
| **BJ** | 16 | 16 | 10 |
| **DV** | 24 | 20 | 20 |
| **EV** | 27 | 25 | 20 |
| **GR** | 41 | 25 | 25 |
| **HW** | 36 | 30 | 27 |
| **KD** | 54 | 25 | 17 |
| **KK** | 14 | 10 | 10 |
| **MT** | 11 | 11 | 11 |
| **MP** | 12 | 12 | 8 |
| **MN** | 14 | 14 | 12 |
| **PV** | 13 | 12 | 9 |
| **Total** | **332** | **242** | **207** |

**Supp. Table 2:** Summary of notes and songs annotated across individuals per location.

| Population | Individual | Songs per indiv | Notes per<br>indv |
| --- | --- | --- | --- |
| ANAIM<br>(N = 13) | AM_01 | 80 | 703 |
|  | AM_02 | 55 | 627 |
|  | AM_03 | 86 | 1347 |
|  | AM_05 | 83 | 1135 |
|  | AM_06 | 80 | 872 |
|  | AM_07 | 80 | 703 |
|  | AM_08 | 83 | 635 |
|  | AM_09 | 90 | 965 |
|  | AM_10 | 63 | 674 |
|  | AM_11 | 68 | 626 |
|  | AM_13 | 100 | 954 |
|  | AM_15 | 77 | 1048 |
|  | AM_17 | 55 | 490 |
| BANDR<br>(N = 14) | BD_02 | 30 | 327 |
|  | BD_03 | 27 | 303 |
|  | BD_07 | 111 | 1001 |
|  | BD_09 | 28 | 238 |
|  | BD_10 | 86 | 943 |
|  | BD_11 | 70 | 656 |
|  | BD_12 | 52 | 567 |
|  | BD_13 | 96 | 1261 |
|  | BD_14 | 96 | 1143 |
|  | BD_15 | 88 | 1026 |
|  | BD_18 | 29 | 296 |
|  | BD_20 | 110 | 1387 |
|  | BD_21 | 67 | 667 |
|  | BD_22 | 110 | 1387 |
| BERJM<br>(N = 13) | BJ_01 | 85 | 905 |
|  | BJ_02 | 85 | 1061 |
|  | BJ_03 | 61 | 694 |
|  | BJ_04 | 86 | 963 |
|  | BJ_05 | 52 | 648 |
|  | BJ_06 | 63 | 623 |
|  | BJ_07 | 88 | 1150 |
|  | BJ_08 | 88 | 1435 |
|  | BJ_10 | 88 | 1072 |
|  | BJ_12 | 87 | 1240 |
|  | BJ_13 | 88 | 1014 |
|  | BJ_14 | 66 | 954 |
|  | BJ_15 | 63 | 543 |
| ERAVI<br>(N = 12) | EV_01 | 20 | 199 |
|  | EV_04 | 27 | 193 |
|  | EV_07 | 94 | 1203 |
|  | EV_08 | 114 | 1492 |
|  | EV_09 | 78 | 966 |
|  | EV_10 | 73 | 947 |
|  | EV_11 | 117 | 1494 |
|  | EV_12 | 52 | 521 |
|  | EV_13 | 142 | 2935 |
|  | EV_14 | 64 | 929 |
|  | EV_15 | 116 | 1709 |
|  | EV_16 | 100 | 1877 |
| GRASS<br>(N = 13) | GR_02 | 87 | 1251 |
|  | GR_03 | 42 | 596 |
|  | GR_04 | 90 | 916 |
|  | GR_05 | 56 | 808 |
|  | GR_06 | 59 | 725 |
|  | GR_07 | 90 | 1098 |
|  | GR_08 | 60 | 570 |
|  | GR_09 | 65 | 913 |
|  | GR_10 | 91 | 1024 |
|  | GR_11 | 85 | 1978 |
|  | GR_12 | 90 | 1075 |
|  | GR_13 | 90 | 1092 |
|  | GR_16 | 95 | 1400 |
| HIGHW<br>(N = 13) | HW_01 | 69 | 747 |
|  | HW_02 | 201 | 2305 |
|  | HW_03 | 114 | 1515 |
|  | HW_04 | 27 | 366 |
|  | HW_05 | 49 | 364 |
|  | HW_06 | 28 | 329 |
|  | HW_07 | 62 | 662 |
|  | HW_08 | 146 | 2536 |
|  | HW_09 | 201 | 3533 |
|  | HW_10 | 29 | 615 |
|  | HW_11 | 22 | 322 |
|  | HW_13 | 24 | 387 |
|  | HW_14 | 28 | 439 |
| KODAI<br>(N = 12) | KD_01 | 24 | 452 |
|  | KD_02 | 90 | 881 |
|  | KD_03 | 90 | 950 |
|  | KD_04 | 90 | 895 |
|  | KD_06 | 90 | 1018 |
|  | KD_07 | 90 | 898 |
|  | KD_08 | 88 | 1146 |
|  | KD_09 | 90 | 1095 |
|  | KD_10 | 90 | 1130 |
|  | KD_11 | 86 | 1008 |
|  | KD_12 | 86 | 936 |
|  | KD_13 | 86 | 866 |
| TOTAL | 90 | 6997 | 87589 |

**Supp. Table 3:** Loadings and proportion of variance explained for the first three principal components (PC1–PC3) from a PCA on eight precipitation-related bioclimatic variables (bio12–bio19) across the sampled locations. PC1 alone accounted for 72.2% of the total variance.

|  | PC1 | PC2 | PC3 |
| --- | --- | --- | --- |
| Variance Explained | 0.722 | 0.157 | 0.058 |
| bio12 | 0.404 | 0.024 | 0.288 |
| bio13 | 0.407 | -0.103 | 0.169 |
| bio14 | 0.361 | 0.142 | -0.575 |
| bio15 | 0.140 | -0.802 | -0.198 |
| bio16 | 0.405 | -0.146 | 0.153 |
| bio17 | 0.328 | 0.424 | -0.478 |
| bio18 | 0.349 | 0.270 | 0.510 |
| bio19 | 0.358 | -0.228 | -0.092 |

**Supp. Table 3a:**
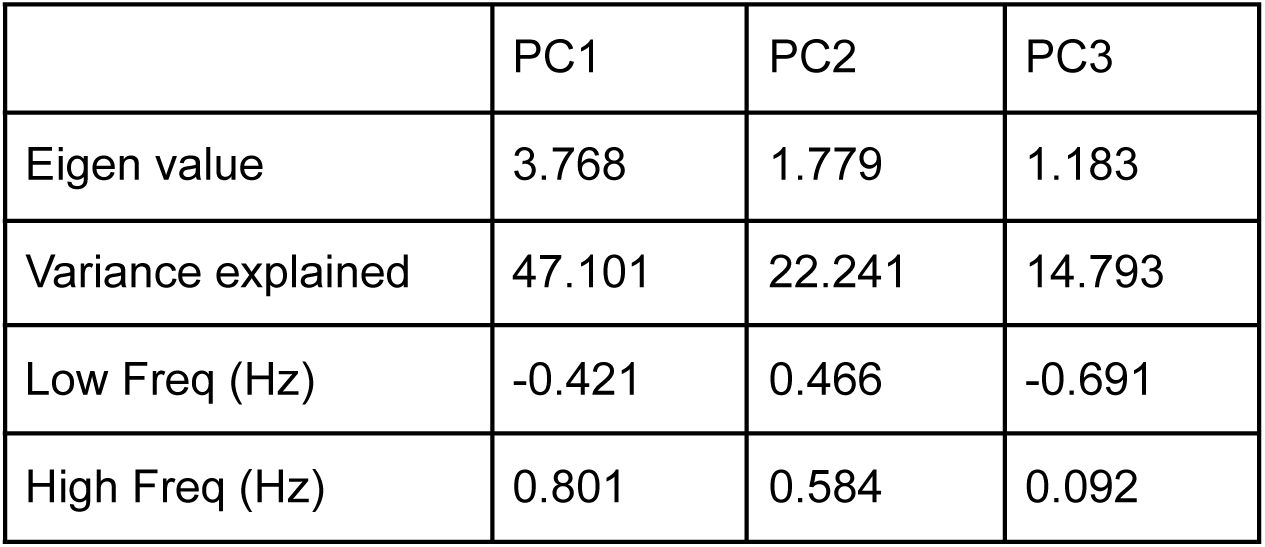
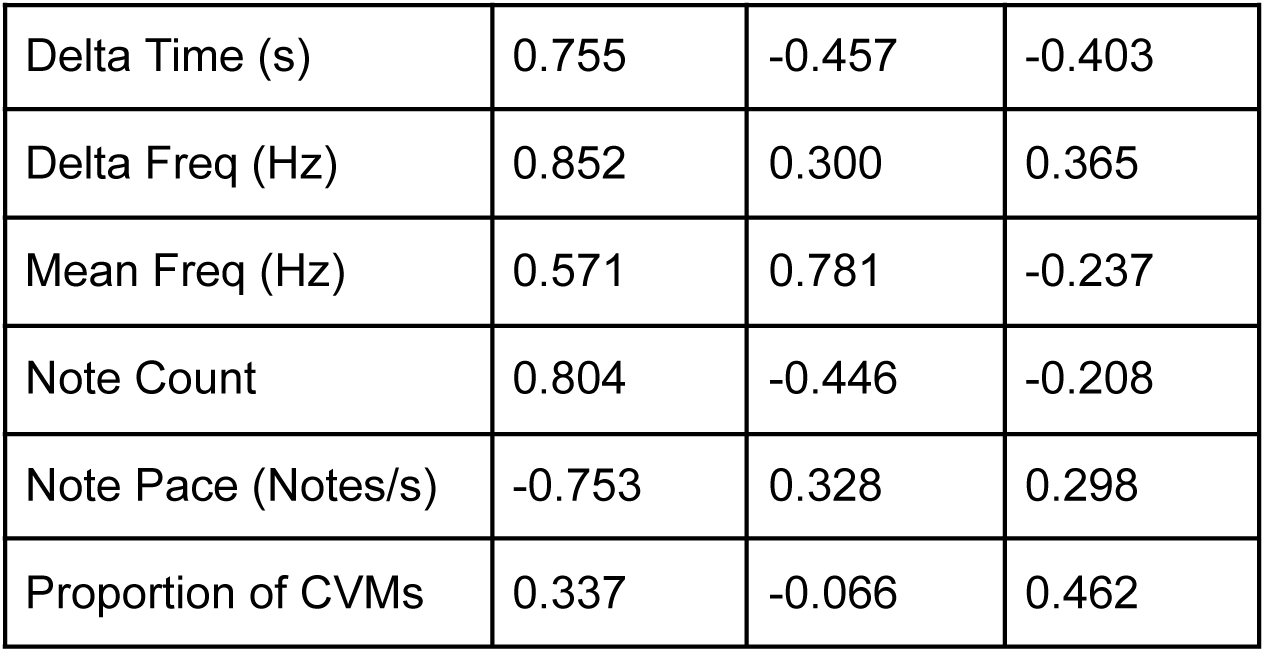
Loadings, eigenvalues, and percentage of variance explained for the first three principal components (PC1–PC3) from a PCA on eight acoustic parameters: spectral features (Low Freq, High Freq, Mean Freq, Delta Freq), temporal features (Delta Time, Note Count, Note Pace), and the proportion of Complex Vocal Mechanisms (CVMs). Together, PC1–PC3 explained 84.1% of the total variance.

**Supp Table 4:**
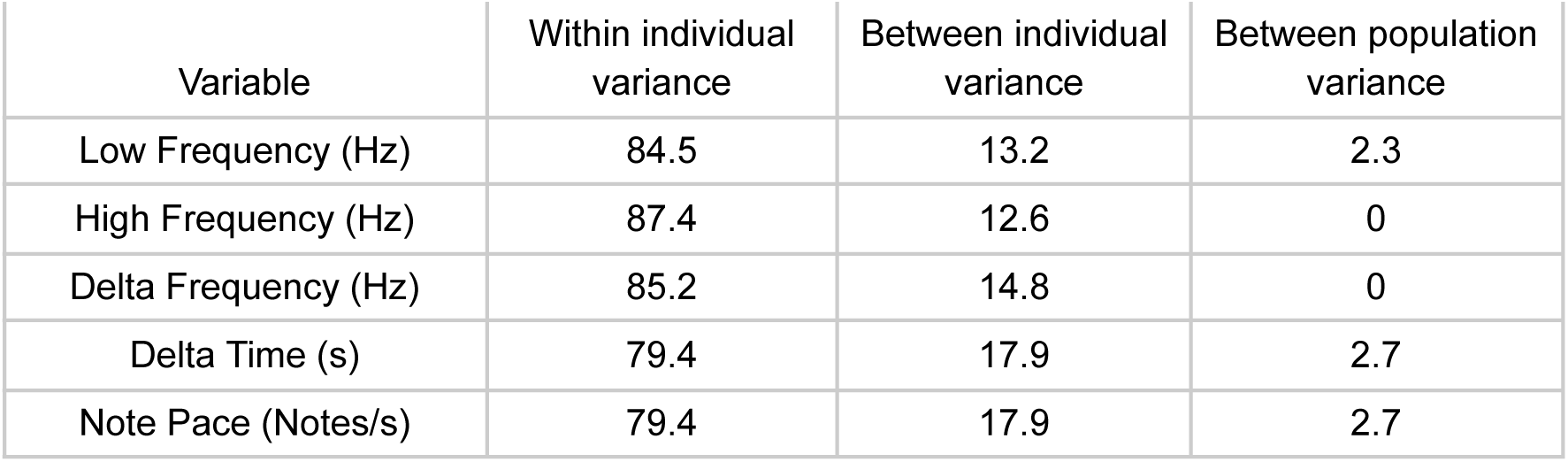
Proportional variance components (%) for song-level acoustic variables, partitioned among three hierarchical levels using linear mixed models with individual nested within population.

**Supp. Table 5:** Variation within and across populations indicated via Permanova results for all levels of N-gram sharing (1-gram note level and 2 - 3g syntax)

| Distance Metric | N-gram-level | DF | PERMANOVA R<sup>2</sup> | PERMANOVA F | PERMANOVA p-value | Betadisp F | Betadisp p |
| --- | --- | --- | --- | --- | --- | --- | --- |
| Jaccard's distance | 1-gram | (6,83) | 0.227 | 4.056 | 0.001 | 3.88 | 0.004 |
| Jaccard's distance | 2-gram | (6,83) | 0.157 | 2.569 | 0.001 | 3.385 | 0.004 |
| Jaccard's distance | 3-gram | (6,83) | 0.157 | 2.575 | 0.001 | **1.714** | **0.16** |
| Distance Metric | n-gram level | DF | PERMANOVA R<sup>2</sup> | PERMANOVA F | PERMANOVA p-value | Betadisp F | Betadisp p |
| Bray–Curtis | 1-gram | (6,83) | 0.165 | 2.731 | 0.001 | 2.394 | 0.045 |
| Bray–Curtis | 2-gram | (6,83) | 0.15 | 2.448 | 0.001 | 2.531 | 0.021 |
| Bray–Curtis | 3-gram | (6,83) | 0.148 | 2.407 | 0.001 | 2.221 | 0.054 |

**Supp. Table 6:** PCA loadings from playback experiment trials conducted.

|  | PC1 | PC2 |
| --- | --- | --- |
| Eigen value | 1.72 | 1.07 |
| Variance explained | 43.08 | 26.67 |
| Average distance to the speaker (m) | -0.35 | **-0.62** |
| Total singing duration (sec) | **0.87** | -0.28 |
| Flights over the speaker | 0.25 | **0.76** |
| Latency to sing (sec) | **-0.89** | 0.18 |

**Supp. Table 8:** Set of tables for full and best fit models generated via Multiple Matrix Regression with Randomisation model. a) Testing genetic distances (FST) and the predicted variable of modes of isolation via IBD, IBE and IBR. b) Testing genetic distances (DPS) and the predicted variable of modes of isolation via IBD, IBE and IBR. c) Results of MMRR models testing Isolation by Distance (IBD), Isolation by Environment (IBE), and Isolation by Resistance (IBR) on 1-, 2-, and 3-gram presence–absence distances. d) *Results of MMRR models testing Isolation by Distance (IBD), Isolation by Environment (IBE), and Isolation by Resistance (IBR) on 1-, 2-, and 3-gram relative abundance distances*.

| Model | Term | Estimate | p | 95% CI Lower | 95% CI Upper | R<sup>2</sup> | F | F p-value |
| --- | --- | --- | --- | --- | --- | --- | --- | --- |
| IBD | IBD | 0.78 | **<0.001** | 0.66 | 0.9 | 0.61 | 118.16 | **<0.001** |
|  | Intercept | 0 | 0.41 | -0.12 | 0.12 |  |  |  |
| IBE | clim_PC1 | -0.15 | 0.47 | -0.34 | 0.04 | 0.07 | 1.98 | 0.54 |
|  | clim_PC2 | 0.23 | 0.24 | 0.04 | 0.41 |  |  |  |
|  | clim_PC3 | -0.07 | 0.68 | -0.25 | 0.12 |  |  |  |
|  | Intercept | 0 | 0.46 | -0.19 | 0.19 |  |  |  |
| IBR | IBR | 0.71 | **<0.001** | 0.57 | 0.84 | 0.5 | 75.59 | **<0.001** |
|  | Intercept | 0 | 0.19 | -0.13 | 0.13 |  |  |  |
| IBD + IBR residuals | IBD | 0.78 | **<0.001** | 0.66 | 0.9 | 0.62 | 61.77 | **<0.001** |
|  | IBR_residuals | 0.12 | 0.4 | 0 | 0.24 |  |  |  |
|  | Intercept | 0 | 0.64 | -0.12 | 0.12 |  |  |  |

b) Testing genetic distances (DPS) and the predicted variable of modes of isolation via IBD, IBE and IBR.
| Model | Term | Estimate | p | 95% CI Lower | 95% CI Upper | R<sup>2</sup> | F | F p-value |
| --- | --- | --- | --- | --- | --- | --- | --- | --- |
| IBD | IBD | 0.21 | 0.46 | 0.02 | 0.4 | 0.04 | 3.51 | 0.46 |
|  | Intercept | 0 | 0.55 | -0.19 | 0.19 |  |  |  |
| IBE | clim_PC1 | -0.29 | 0.15 | -0.48 | -0.11 | 0.1 | 2.64 | 0.45 |
|  | clim_PC2 | -0.09 | 0.62 | -0.28 | 0.09 |  |  |  |
|  | clim_PC3 | 0 | 0.98 | -0.18 | 0.19 |  |  |  |
|  | Intercept | 0 | 0.53 | -0.18 | 0.18 |  |  |  |
| IBR | IBR | 0.16 | 0.63 | -0.03 | 0.35 | 0.03 | 2 | 0.63 |
|  | Intercept | 0 | 0.57 | -0.19 | 0.19 |  |  |  |

| N-gram | Model | Term | Estimate | p | 95% CI Lower | 95% CI Upper | R<sup>2</sup> | F | F p-value |
| --- | --- | --- | --- | --- | --- | --- | --- | --- | --- |
| 1-gram | IBD | IBD | 0.07 | 0.84 | -0.32 | 0.47 | 0.01 | 0.1 | 0.84 |
|  |  | Intercept | 0 | 0.64 | -0.39 | 0.39 |  |  |  |
|  | IBE | clim_PC1 | 0.31 | 0.14 | -0.04 | 0.65 | 0.38 | 3.47 | 0.1 |
|  |  | clim_PC2 | -0.08 | 0.75 | -0.42 | 0.26 |  |  |  |
|  |  | clim_PC3 | 0.44 | 0.09 | 0.09 | 0.8 |  |  |  |
|  |  | Intercept | 0 | 0.11 | -0.32 | 0.32 |  |  |  |
|  | IBR | IBR | -0.15 | 0.86 | -0.54 | 0.25 | 0.02 | 0.41 | 0.86 |
|  |  | Intercept | 0 | 0.32 | -0.38 | 0.38 |  |  |  |
| 2-gram | IBD | IBD | -0.31 | 0.28 | -0.69 | 0.07 | 0.1 | 2.01 | 0.28 |
|  |  | Intercept | 0 | 0.61 | -0.37 | 0.37 |  |  |  |
|  | IBE | clim_PC1 | 0.14 | 0.66 | -0.26 | 0.54 | 0.17 | 1.15 | 0.45 |
|  |  | clim_PC2 | -0.4 | 0.16 | -0.8 | -0.01 |  |  |  |
|  |  | clim_PC3 | -0.03 | 0.9 | -0.44 | 0.38 |  |  |  |
|  |  | Intercept | 0 | 0.76 | -0.38 | 0.38 |  |  |  |
|  | IBR | IBR | -0.31 | 0.22 | -0.68 | 0.07 | 0.09 | 1.96 | 0.22 |
|  |  | Intercept | 0 | 0.93 | -0.37 | 0.37 |  |  |  |
| 3-gram | IBD | IBD | -0.08 | 0.77 | -0.48 | 0.31 | 0.01 | 0.14 | 0.77 |
|  |  | Intercept | 0 | 0.31 | -0.39 | 0.39 |  |  |  |
|  | IBE | **clim_PC1** | **0.68** | **0.01** | **0.39** | **0.98** | **0.55** | **7** | **0.01** |
|  |  | **clim_PC2** | **-0.35** | **0.05** | **-0.64** | **-0.06** |  |  |  |
|  |  | clim_PC3 | -0.02 | 0.92 | -0.32 | 0.28 |  |  |  |
|  |  | Intercept | 0 | 0.02 | -0.28 | 0.28 |  |  |  |
|  | IBR | IBR | -0.19 | 0.54 | -0.58 | 0.2 | 0.03 | 0.68 | 0.54 |
|  |  | Intercept | 0 | 0.23 | -0.38 | 0.38 |  |  |  |

| N-gram | Model | Term | Estimate | p | 95% CI Lower | 95% CI Upper | R<sup>2</sup> | F | F p-value |
| --- | --- | --- | --- | --- | --- | --- | --- | --- | --- |
| 1-gram | IBD | IBD | -0.04 | 0.92 | -0.43 | 0.36 | 0 | 0.02 | 0.92 |
|  |  | Intercept | 0 | 0.78 | -0.39 | 0.39 |  |  |  |
|  | IBE | clim_PC1 | 0.12 | 0.56 | -0.24 | 0.47 | 0.35 | 3.09 | 0.1 |
|  |  | clim_PC2 | -0.21 | 0.32 | -0.57 | 0.14 |  |  |  |
|  |  | clim_PC3 | 0.47 | 0.07 | 0.11 | 0.83 |  |  |  |
|  |  | Intercept | 0 | 0.73 | -0.33 | 0.33 |  |  |  |
|  | IBR | IBR | -0.33 | 0.28 | -0.7 | 0.05 | 0.11 | 2.29 | 0.28 |
|  |  | Intercept | 0 | 0.51 | -0.37 | 0.37 |  |  |  |
| 2-gram | IBD | IBD | 0.04 | 0.93 | -0.35 | 0.44 | 0 | 0.03 | 0.93 |
|  |  | Intercept | 0 | 0.18 | -0.39 | 0.39 |  |  |  |
|  | IBE | clim_PC1 | 0.02 | 0.92 | -0.37 | 0.41 | 0.2 | 1.38 | 0.46 |
|  |  | clim_PC2 | 0.17 | 0.55 | -0.22 | 0.56 |  |  |  |
|  |  | clim_PC3 | 0.44 | 0.16 | 0.04 | 0.84 |  |  |  |
|  |  | Intercept | 0 | 0.18 | -0.37 | 0.37 |  |  |  |
|  | IBR | IBR | 0.04 | 0.94 | -0.36 | 0.44 | 0 | 0.03 | 0.94 |
|  |  | Intercept | 0 | 0.1 | -0.39 | 0.39 |  |  |  |
| 3-gram | IBD | IBD | 0.1 | 0.55 | -0.29 | 0.5 | 0.01 | 0.2 | 0.55 |
|  |  | Intercept | 0 | 0.67 | -0.39 | 0.39 |  |  |  |
|  | IBE | clim_PC1 | 0.2 | 0.46 | -0.23 | 0.63 | 0.05 | 0.31 | 0.79 |
|  |  | clim_PC2 | -0.06 | 0.79 | -0.49 | 0.36 |  |  |  |
|  |  | clim_PC3 | 0.05 | 0.82 | -0.39 | 0.49 |  |  |  |
|  |  | Intercept | 0 | 0.27 | -0.4 | 0.4 |  |  |  |
|  | IBR | IBR | -0.06 | 0.98 | -0.45 | 0.34 | 0 | 0.06 | 0.98 |
|  |  | Intercept | 0 | 0.43 | -0.39 | 0.39 |  |  |  |

**Supp. Table 9:** Partial Mantel test results of song distances and genetic distances (FST) accounting for geographic distances.

|  | **DPS** |  | **FST** |  |
| --- | --- | --- | --- | --- |
| **Song distances -**<br>**Presence Absence** | **R stat** | **p-value** | **R stat** | **p-value** |
| **1-gram** | -0.4583 | 0.959 | -0.49 | 0.95 |
| **2-gram** | -0.4097 | 0.863 | -0.22 | 0.79 |
| **3-gram** | -0.5464 | 1 | -0.44 | 0.99 |
| **Song distances -**<br>**Relative abundances** | **R stat** | **p-value** | **R stat** | **p-value** |
| **1-gram** | -0.0724 | 0.615 | -0.41 | 0.92 |
| **2-gram** | -0.3987 | 0.873 | -0.17 | 0.70 |
| **3-gram** | -0.1739 | 0.884 | -0.35 | 1.00 |

