## Supplementary Methods for "Songs show strong individual-level variation and weak population-level dialects in a tropical songbird with genetic structure"

*S.1.0 Acoustic data analysis:*

S1.1) Annotations and spectral data analysis, along with Complex Vocal Mechanisms (CVMs)

All songs were manually annotated by CA at the note level in Raven Pro v1.6 (K. Lisa Yang Centre for Conservation Bioacoustics at the Cornell Lab of Ornithology) by generating spectrograms with a 512-sample Hann window, 50% overlap, and a 512 DFT. A single note was judged based on its separation from the following consecutive note by any space of approximately >0.02s - a natural breath unit of small passerines used by some authors (Singh & Price, 2015). Notes separated by < 0.02s were marked as a part of a Complex Vocal Mechanism (CVM) (Purushotham & Robin, 2016), possibly from a non-linear phenomenon or two-voice complexity mechanism of sound production using both sides of the syrinx (Zollinger et al., 2008). Overlapping notes falling under a CVM were cross-checked (by CA and AGP) to ensure that they were annotated independently of each other. We extracted the following spectral variables at the song level: low frequency (Hz), high frequency (Hz), delta time (s), delta frequency (Hz), mean frequency (Hz), note count (number of notes in a song), note pace (note count/delta time(sec)) and proportion of Complex Vocal Mechanisms (CVMs) in a song (instances of CVMs in a song/ note count).

S1.2) Syntax assessment - note classification for syllable extraction

First, we extracted notes using time- and frequency-level note parameters from the manual annotations (described above), denoised them, and saved each note as a separate WAV file. All analyses were conducted in Python (version 3.10.2) using the following packages: NumPy, pandas, SciPy (signal processing module) and librosa. Next, we estimated spectrogram cross-correlation (SPCC) scores by comparing each test note iteratively with a manually classified library of 1,000 notes (Sawant et al., 2025). Each test note was compared to all notes in the library using spectrogram cross-correlation to obtain a score. If any test note had a score > 0.9925 (high confidence threshold) with a library note, it was immediately assigned that note's class, and the loop stopped. If no correlation exceeded 0.9925 but at least one score fell between 0.986 and 0.9925, the test note was assigned the class of the best-matching library note in that range. If the maximum score was < 0.986 (low confidence threshold), the note was treated as a new note. All analyses were conducted in Python using the following libraries: NumPy, pandas, SciPy (signal processing module) and librosa. Thresholds were determined using Receiver Operating Characteristic (ROC) optimisation. This process was parallelised to speed up the classification. We manually validated the classification by randomly selecting 100 notes from each of the seven populations, achieving a classification accuracy of ~70%.

S1.3) Inferring syllable sharing within and across populations using N-gram analysis

Most studies examine cultural diversity in terms of repertoire size (Fayet et al., 2014; Pérez-Granados et al., 2016; Sebastián-González et al., 2025; Sprau & Mundry, 2010; J. M. Williams & Slater, 1990); here, we estimate repertoire size and also utilise a linguistic tool, N-grams, to identify syntax by analysing the ordering of notes to form syllables (Sawant et al., 2025).  We visualised song sharing at both the note (1-gram) and syllable (2-gram and 3-gram) levels using classified notes (from Section 1.2) across all 90 individuals (custom script). To minimise the impact of automated classification errors and random variation, we filtered for occurrences at each gram level: 1-grams with more than 5 per population, and 2-grams and 3-grams with more than 2 times per population. We then visualised the sharing of each N-gram type within and across populations (Venn diagrams) and examined patterns of increase or decrease in N-gram sharing across all seven populations, as well as N-grams unique to each population.

*S2.0) Playback experiment*

S2.1) Playback experiment data collection

We gathered GPS coordinates for playback trials conducted on all unringed individuals, enabling subsequent trials to be conducted at the exact locations. Trials were conducted over two seasons, winter (December 2024) and summer (March 2025). The trials consisted of two sessions per day, held in the morning (6:00–9:00 am) and the evening (3:00–6:00 pm). It was ensured that the same individual received only one trial per session and that neighbouring birds were not tested with the same stimulus on the same day (Lipshutz et al., 2017). We used a dual-observer system in which both observers vocally annotated behaviours into a microphone, with at least one observer maintaining a line of sight with the focal bird as it responded to the playback. Trials were performed simultaneously across non-adjacent focal individuals by three teams (each with 2 people) and out of playback earshot, with observers shuffled between sessions and blind to trial identity.

All sound files were filtered (using a bandpass filter with a frequency range of 1–10 kHz) and normalised (using Ocenaudio) before being broadcast at 65 dB SPL through a wireless speaker (JBL Go 3, connected via Bluetooth to a smartphone) placed 1m above the ground in the centre of the territory. If needed, birds were briefly lured into view with the white-bellied sholakili whistle call on Merlin (10s clip - on repeat for one minute) before the trial began. Once the bird was spotted, we began the playback trial after a one-minute silence. Data from incomplete or interrupted trials (< 6 minutes) were excluded from the analyses (n = 3 ), resulting in a final pool of 136 trials.

S2.2) Playback experiment data analysis

Experiments were transcribed, blind to the recordings, into 10-second intervals in an ethogram by manually listening to the recordings (RS). We ran two models with vocal and physical responses as the response variables, using the first two principal components and their loadings. The predictor variables included the population from which the playback songs originated, and the random effects included were the observer annotating the experiment and the focal bird.

S3.0) Genomic data - ddRADseq library preparation

CA and NG prepared an initial 159-sample ddRADseq library following the modified protocol of Tyagi et al. (2024), as detailed in Praveen et al. (2024). An additional 83 samples were sequenced and demultiplexed at an external facility (Strand Life Sciences Pvt Ltd, Bangalore) using the same set of restriction enzymes. All libraries were sequenced on an Illumina Novaseq 6000 platform. Briefly, all samples were digested with two restriction enzymes: SphI (a 6-bp cutter) and MluCI (a 4-bp cutter). Digested samples were then indexed with custom (dual) adapters (Tyagi et al, 2024). Size selection was done using Ampure XP beads, and library quality was assessed using an Agilent TapeStation. Libraries were then pooled at equimolar ratios and sequenced on an Illumina NovaSeq 6000 (150 bp x 2) in two different runs. In addition to the initial libraries, 83 more samples were sequenced and demultiplexed at an external facility (Strand Life Sciences Pvt Ltd, Bangalore) in two batches using the same set of restriction enzymes. Reads from all four runs were merged and processed for further analysis.

S3.1) Genomic data processing - SNP genotyping

We assessed the quality of the demultiplexed reads using FastQC (Andrews, 2010) and trimmed the reads for adapters and poor-quality bases using Trimmomatic v0.39 (Bolger et al., 2014). An interactive assembly pipeline, ipyrad v.0.9.105 (Eaton & Overcast, 2020), was used for variant calling using a reference-assembly Method, mapping reads to a high-quality reference genome of the White-bellied Sholakili (Vinay et al., 2025). Specifically, we filtered the reads for quality by converting bases with Phred scores below 20 to N, and any reads with more than 5 Ns were discarded. Furthermore, any reads with post-quality filtering below 35 bp in length were also discarded. The minimum depth for statistical and majority-rule base calling was set to 5, with remaining parameters set to their default values. For variant filtering, we first removed individuals with > 50% missing data (N = 18). We also examined kinship using NGSrelate (Korneliussen & Moltke, 2015) and excluded related individuals with Rab scores > 0.1 (N = 17). We then filtered for bi-allelic SNPs (--min-alleles 2 and –max-alleles 2), used a minimum mean site depth of eight across all individuals (--min-meanDP 8) and a maximum mean site depth of 25 across all individuals (--max-meanDP 25). We then excluded sites with more than 25% missing data (--max-missing 0.75) and out of Hardy-Weinberg equilibrium (--hwe 0.05), and retained sites with a minor allele frequency greater than 0.05 (--maf 0.05). To account for linkage disequilibrium (only for PCA and ADMIXTURE analyses), we thinned the data to 10,000 bp (--thin 10000). All filtering was performed using VCFTools v0.1.13 (Auton and Marcketta 2009).

S3.2) Coancestry and estimated effective migration surfaces (EEMS)

We also assessed genetic co-ancestry among individuals within and across locations (using a haplotype-based population inference approach implemented in fineRADstructure v.0.1), which derives a coancestry matrix from the SNP dataset using the same filtering criteria described for the SNP dataset, without thinning. Analyses were performed using default settings. The results were visualised using the R scripts provided with the software package. To detect barriers to dispersal that may be an alternative to isolation-by-distance, we employed FEEMS (Fast Estimation of Effective Migration Surfaces). This technique generates migration surfaces and quantifies pairwise edge estimates between nodes, representing effective migration rates that deviate from isolation-by-distance (IBD) expectations. Cross-validation was performed over lambda values ranging from 0.001 to 24, with the optimal lambda for the spatial graph selected by minimising cross-validation error.
